# Signaling downstream of tumor-stroma interaction regulates mucinous colorectal adenocarcinoma apicobasal polarity

**DOI:** 10.1101/2024.11.20.624298

**Authors:** Nicolas Pasquier, Aleksi Isomursu, Hellyeh Hamidi, Jacques R.R. Mathieu, Jouni Härkönen, Gautier Follain, Christophe Desterke, Zoé Fusilier, Junel Solis, Irina Belaya, Pasi Kankaanpää, Valeria Barresi, Jérôme Cartry, Sabrina Bedja, Fanny Jaulin, Johanna Ivaska

## Abstract

Mucinous colorectal carcinoma (MUC CRC) dissemination into the tumor stroma and metastasis to multiple organs, including the peritoneum, is associated with poor prognosis. Disseminating MUC CRCs exhibit either a conventional ‘apical-in’ or an inverted ‘apical-out’ polarity phenotype that influence patient outcome. Identifying the mechanisms controlling MUC CRC polarity is critical to understand disease progression. Here, we analyze patient-derived MUC CRC xenografts, with apical-in or apical-out polarity, ex vivo or within collagen gels to mimic the peritumoral stroma. Single-cell analyses reveal α2β1-integrin as a key collagen-binding receptor in these models. Collagen–α2β1-integrin interaction activates Src and ERK/MAPK signaling and upregulates the expression of SorLA, an endosomal sorting receptor. SorLA supports apical-in polarity and carcinoma-stroma interactions by promoting integrin recycling to the plasma membrane and HER2/HER3 expression through a positive feedback mechanism. Accordingly, we observe positive correlation between HER2, HER3 and SorLA in patient samples with the highest HER2 expression in apical-in-presenting tissues. Treatment of tumor spheres with clinically relevant HER2/HER3-targeting antibodies reverts sphere polarity and impedes collagen remodeling and adhesion to mouse peritoneum. This SorLA—integrin—HER2/HER3 signaling axis may represent a basis for MUC CRC-patient stratification and shed light on other carcinomas with similar apical-out phenotypes.

## Introduction

Apicobasal polarization is a critical cellular process that is essential for normal tissue function. Its disruption is implicated in various pathological conditions including cancer (Macara and McCaffrey, 2013; Wodarz & Näthke 2007; Onuma & Inoue 2022). While the loss of polarity has been extensively studied, the phenomenon of inverted polarity (also referred to as apical-out polarity), where the apical and basolateral domains are reversed, is less understood.

Apical-out topology, characterized by the mislocalization of apical and basolateral markers (Lee and Vasioukhin, 2008; Peglion et al., 2023, Pasquier et al., 2024, Verdú et al., 2011), is a hallmark of several aggressive cancer subtypes (Jakubowska et al., 2016), including micropapillary carcinomas and mucinous colorectal adenocarcinomas (MUC CRC). Micropapillary carcinoma exhibit a fully inverted apicobasal polarity (Verras et al., 2022). This type of cancer is diagnosed upon detection of an apical-out polarity pattern. MUC CRC represents a distinct subtype of colorectal carcinoma (CRC), characterized by abundant mucinous components constituting at least 50% of the tumor volume. CRC is the third most common cancer globally and the second leading cause of cancer-related deaths (Siegel et al., 2023). MUC CRC constitutes 10-15% of CRC cases, predominantly affecting young women and the right colon. It carries a poor prognosis, particularly in metastatic disease, with a high incidence of peritoneal metastasis (Mekenkamp et al., 2012; Luo et al., 2019; Bettington et al., 2013; Yamane et al., 2014; Libanje et al., 2019; Hugen et al., 2014). Therefore, there is an urgent need to better understand the mechanisms contributing to acquisition of inverted polarity in cancer.

Histological analyses have revealed that MUC CRCs disseminate not as individual cells but rather as clusters of hundreds of cells called tumor spheres with inverted polarity (TSIPs) (Zajac et al., 2018). These structures, first identified in patients’ peritoneal effusions, act as tumor intermediates arising from the primary tumor and invading tissues to form metastasis in the peritoneum, facilitating cancer motility and dissemination (Pagès et al., 2022).

While these TSIPs maintain an inverted (apical-out) polarity in suspension, they display distinct responses within tumor stroma. Once in contact with the extracellular matrix (ECM), TSIPs either revert to a normal (apical-in) polarity or remain inverted (apical-out). The mechanisms underpinning these differential polarity responses to the stromal environment are not fully understood, although the TGF-β pathway has been implicated (Okuyama et al., 2016, Onuma et al., 2021, Canet-Jourdan et al., 2022). Apical-out polarity phenotype has recently been shown to have clinically relevant manifestations. It is supportive of a collective amoeboid mode of migration enhancing tumor invasion efficiency (Pagès et al., 2022), and significantly impacts tumor sensitivity to anti-cancer drugs (Ashley et al, 2019).

There is growing evidence from different cancer types that tumor-ECM interactions regulate cancer progression in numerous ways (Hamidi et al., 2018; Rafaeva & Erler, 2020). Cancer cell-ECM interactions and the mechanical properties of the ECM, orchestrated strongly by cancer associated fibroblasts (Pickup et al., 2014; Calvo et al., 2013) and cancer cells via matrix remodeling, regulate cancer cell proliferation, migration and invasion through integrin-mediated signaling and integrin and actin mediated force transduction to the nucleus (Kechagia et al., 2019). While integrins are well-established mediators of mechanotransduction especially in invasive cancer, their role in regulation of cancer cell polarity and how this may be regulated by endosomal receptor traffic is less well understood and has only recently been investigated in prostate cancer (Román-Fernández et al., 2023).

Here, using patient-derived MUC CRC tumor spheres, we uncovered a novel mechanism of TSIP polarity orientation. Tumor sphere contact with stromal collagen induces α2β1-integrin signaling to Src and ERK/MAPK, upregulates the HER2/HER3/SorLA complex activity and increases SorLA-dependent β1-integrin recycling to the plasma membrane. This maintains tumor–stroma adhesion to support apical-in polarity. Increased ligand-induced HER3 signaling is sufficient to trigger apical-in topology in inherently apical-out tumors. Conversely, clinically used HER2/HER3 targeting antibodies inhibit apical-in phenotype. In clinical specimens, SorLA/HER2/HER3 levels are positively correlated and highest in apical-in tumors. Our study demonstrates the central role of SorLA in the regulation of integrin and HER-family receptor function in CRC, significantly contributing to our understanding of polarity regulation in this aggressive cancer type and paving the way for further investigations into the relevance of this signaling axis in MUC CRC progression. Moreover, as inverted polarity is characteristic to a number of aggressive cancers, these findings have broader implications beyond MUC CRC.

## Results

### Mucinous CRC polarity is regulated by the cells’ proximity to the ECM

To investigate the molecular mechanisms of ECM-regulated MUC CRC polarity we used previously described patient-derived xenograft (PDX) models. We focused on three PDXs from the CReMEC bank (Julien et al. 2012) representing mucinous histotypes of CRCs (Canet-Jourdan et al. 2022) and derived either from a peritoneal metastasis (PDX#1, for LRB-009C) or from the primary tumor (PDX#2, for IGR-012P; PDX#3, for IGR-014P) (see Table 1). We cultured the MUC CRC PDX-derived cell clusters (from here on referred to as tumoroids) either in suspension (three days) or in suspension followed by embedding in collagen-I gels, a surrogate for the cancer stroma (Wolf & Friedl, 2011). These two setups mimic the *in vivo* relevant scenarios of MUC CRC progression as peritoneal effusions or as cell clusters in tissue, respectively. In order to quantitatively score polarity phenotypes, we computed a polarity score according to three parameters: presence/absence of a lumen, presence/absence of protrusions and Ezrin fluorescence ratio between the cortical and luminal sides (Fig. S1A). Confocal microscopy imaging of the apical marker Ezrin and quantifications revealed that two of the three PDX tumoroids (PDX#1 and PDX#2) maintain apical-out polarity in both suspension and collagen, whereas PDX#3 tumoroids revert to an apical-in polarity in collagen, forming Ezrin-positive lumens (Fig. 1A,B). These data are concordant with previous studies indicating that 66% of patient-derived TSIPs retain their apico-basolateral polarity in suspension and in collagen (Zajac et al., 2018; Canet-Jourdan et al., 2022). Characteristically, MUC CRCs secrete high levels of free mucin. Upon staining with WGA-lectin, we were able to visualize mucin secretions by the tumoroids. In the non-ECM responsive PDX#1 and PDX#2, mucin surrounded the spheres, indicative of inverted polarity while in PDX#3 the mucin was concentrated within the lumen confirming the polarity reversion–from apical-out to apical-in–in this model (Fig. S1B). To investigate the collagen-induced polarity reversal in detail, we embedded the PDX#3 tumoroids into collagen gels containing fluorescently labelled collagen and performed live-imaging using a filamentous actin live-cell dye (SiR-Actin). We observed dynamic contact between tumoroid cells and the ECM and clear formation of lumen-like structures and protrusions as early as 5 hours post-embedding (Fig. 1C, Video 1). Together, these data imply that only the PDX#3 tumoroids are able to respond to their ECM environment.

**FIG1.**
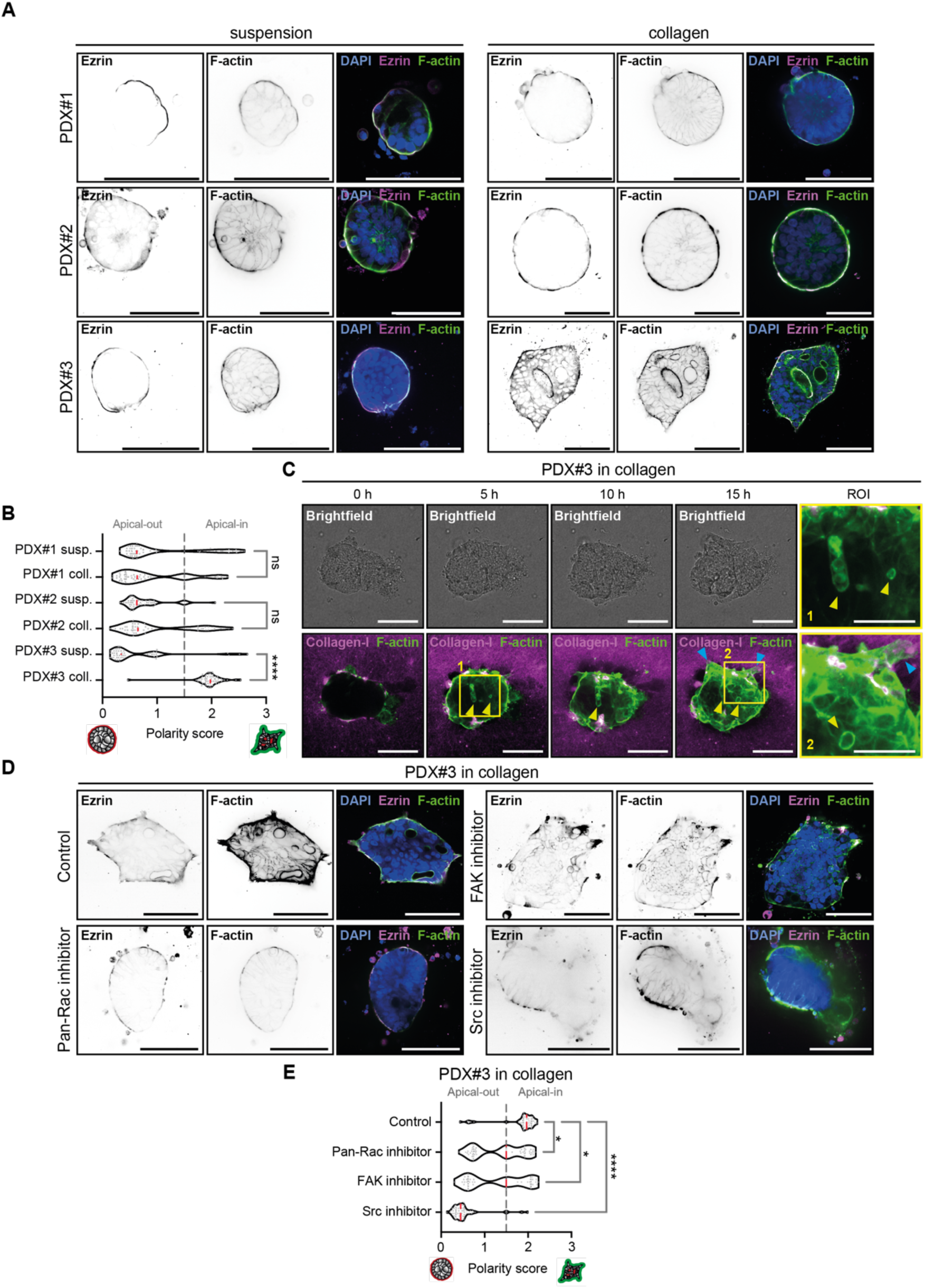
Apicobasal polarity of mucinous CRC PDXs is controlled by the focal adhesion signaling pathway. (A, B) Representative immunofluorescence images of PDX#1-, PDX#2- and PDX#3-generated tumoroids, fixed in suspension or embedded in collagen and stained for Ezrin (polarity marker), F-actin and DAPI (A). Comparison of polarity scores amongst the models is shown (B) [n_PDX#1_ = 51 (suspension), 49 (collagen); n_PDX#2_ = 43 (suspension), 51 (collagen); n_PDX#3_ = 51 (suspension), 52 (collagen); Kruskal-Wallis test with Dunn’s multiple comparison test, p-value_PDX#1_>0.9999, p-value_PDX#2_>0.9999, p-value_PDX#3_<0.0001]. (C) Representative snapshots of live-imaging of PDX#3 tumoroid polarity reversion following embedding in fluorescently labelled collagen (time point 0) over 15 h. Tumoroids were stained with SiR-actin for visualization. Yellow arrows indicate lumen-like structures, blue arrows indicate protrusions. (D, E) Representative immunofluorescence images of PDX#3 tumoroids in collagen treated with either a Pan-Rac inhibitor (EHT-1864, 5 μM), a FAK inhibitor (FAK14, 10 μM) or a Src inhibitor (saracatinib, 1 μM) and stained for Ezrin (D). Analysis of polarity scores after the treatments is shown (E) [n = 45 (control); 53 (EHT-1864); 36 (FAK14) and 35 (saracatinib) PDX#3 tumoroids; Kruskal-Wallis test with Dunn’s multiple comparison test; p-value = 0.0224 (EHT-1864), 0.0204 (FAK14), < 0.0001 (saracatinib)]. Scale bars, 100 μm (main), 50 μm (insets).

**Table 1.** – MUC CRC PDXs used in the study.

| Models | CRemeC ID | Origin | Profiling |
| --- | --- | --- | --- |
| PDX#1 | LRB-009C | Metastatic (peritoneum) | MSS, BRAF WT, KRAS<sup>G12V</sup> |
| PDX#2 | IGR-012P | Primary | MSS, BRAF WT, KRAS<sup>G13D</sup> |
| PDX#3 | IGR-014P | Primary | MSS, BRAF WT, KRAS<sup>G12D</sup> |

### Focal adhesion pathway signaling regulates collagen-induced polarity reversion

To understand the differential responses of the three tumoroid models to collagen embedding, we performed pathway analysis on available transcriptome data (GSE152299) for the three PDX models (suspension versus collagen-embedded tumoroids) (Canet-Jourdan et al., 2022). These analyses revealed the Focal Adhesion Pathway as a top upregulated pathway in the matrix-responsive PDX#3 tumoroids following embedding in collagen. Upon plotting the 45 top differentially regulated genes in the KEGG Focal Adhesion Pathway, we noticed a clear distinction between PDX#3 gene expression in collagen compared to the PDX#1 and PDX#2 tumoroids (Fig. S1C). The corresponding GSEA enrichment plot (Canet-Jourdan et al. 2022) as well as the Principal Component Analysis (PCA) based on the KEGG Focal Adhesion geneset confirmed a clear shift in the transcriptome signature of PDX#3 tumoroids upon collagen embedding (Fig. S1D). These data prompted us to test the functional relevance of the key focal adhesion pathway signaling components on the collagen-induced polarity reversal. We inhibited the small GTPase Rac and the tyrosine kinases focal adhesion kinase (FAK) and Src with established inhibitors (Onesto et al., 2008; Hochwald et al., 2009; Kawata et al., 2022). Each of these inhibitors dampened the polarity shift of PDX#3 in collagen, reducing protrusions, the number of lumens and the luminal/cortical Ezrin signal ratio (Fig 1D-E). Concordantly, Src phosphorylation was induced in PDX#3 upon embedding in collagen (Fig S1E-F), indicating that cell-ECM interaction triggers focal adhesion pathway signaling and contributes to PDX#3 polarity reversal in collagen.

### Collagen-binding integrins are upstream of the polarity establishment signaling pathway in CRC tumoroids

Integrins, in particular β1-integrin-containing heterodimers, are the principal receptors for the ECM upstream of the focal adhesion pathway and represent established polarity markers in epithelia (Manninen 2015; Zajac et al., 2018). We visualized β1-integrin localization and activity in the tumoroids (suspension and collagen). In PDX#3 tumoroids, the localization of β1-integrins (detected with the P5D2 antibody) shifted from lateral (cell-cell) to basal (cell-ECM) and the amount of active β1 (detected with the 12G10 antibody) increased upon contact with the ECM (Fig. 2A). In contrast, β1-integrin intensity was lower in the PDX#1 and PDX#2 tumoroids and the receptor’s localization was not clearly altered despite the cells being embedded in collagen (Fig. 2A).

**FIG2.**
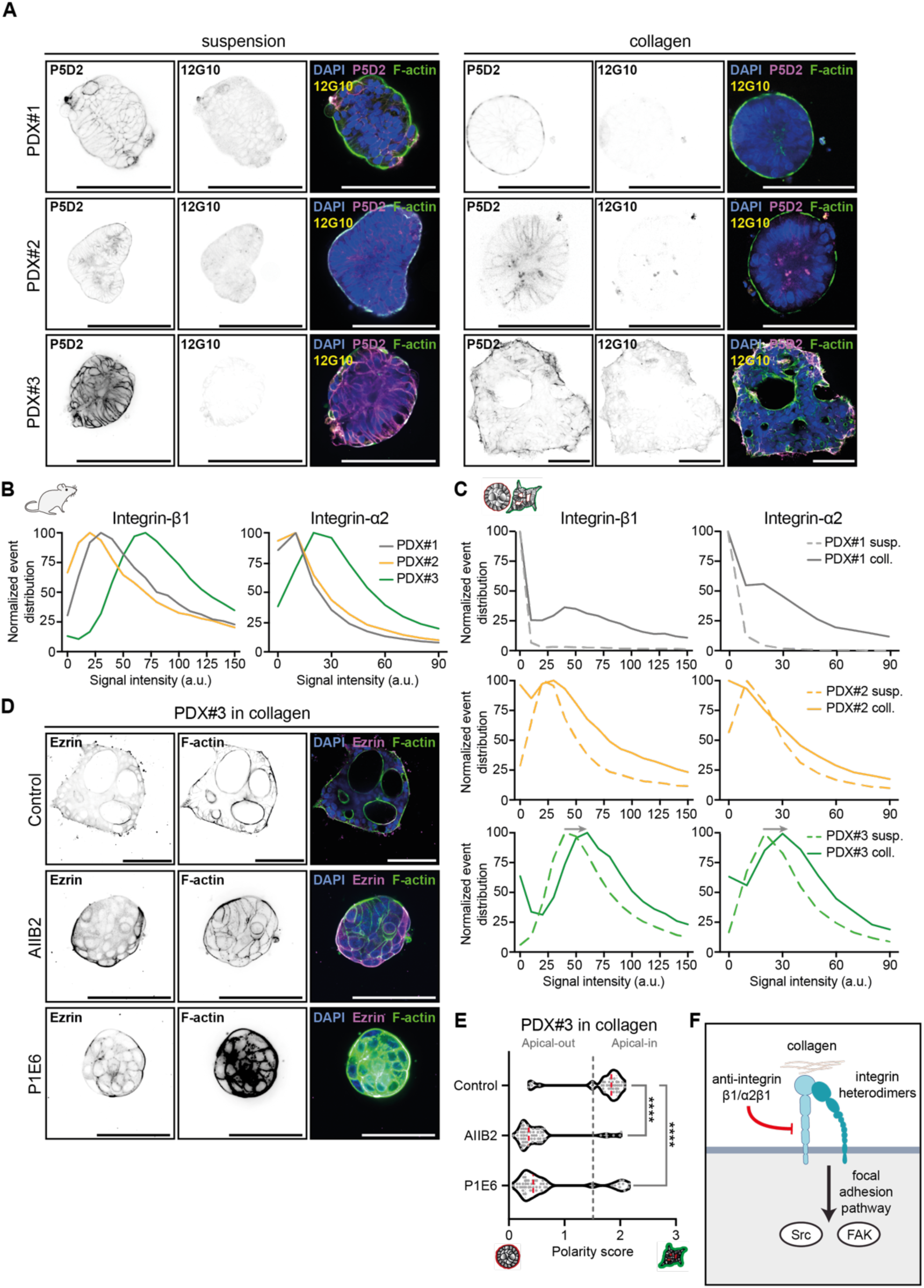
Surface expression of integrin heterodimer α_2_β_1_ varies among PDX models and matrix conditions. (A) Representative immunofluorescence images of PDX#1-, PDX#2- and PDX#3-generated tumoroids in suspension and in collagen, stained for total integrin-β1 (P5D2), active integrin-β1 (12G10), F-actin and DAPI. (B) Distribution of the integrin-β1 and integrin-α2 surface expression signal intensity in PDX#1, PDX#2 and PDX#3 tumors out of mice (cells > 36000). (C) Distribution of integrin-β1 and integrin-α2 surface expression signal intensity in PDX#1-, PDX#2- and PDX#3-generated tumoroids in suspension and in collagen (cells > 7000). (D) Representative immunofluorescence images of PDX#1-, PDX#2- and PDX#3-generated tumoroids in suspension and in collagen treated with an integrin-β1 blocking antibody (AIIB2, 1 μg/ml) or with an integrin-α2 blocking antibody (P1E6, 10 μg/ml) and stained for Ezrin, F-actin and nuclei (DAPI). (E) Comparison of polarity scores of PDX#3 in collagen after integrin-β1 and integrin-α2 blocking [(n = 40 (Control); 61 (AIIB2); 51 (P1E6); Kruskal-Wallis test with Dunn’s multiple comparison test; p-value <0.0001 (AIIB2 and for P1E6)]. (F) Proposed mechanism of collagen-induced signaling and activation of Src and FAK pathways in MUC CRCs. Scale bars, 100 μm.

To assess which β1-integrin heterodimers are involved, we applied high-dimensional mass cytometry analysis, focusing on cell-surface integrin receptor expression at the single cell resolution in the PDX models under different conditions. We designed an extensive antibody panel including all the human ECM-binding integrins (see Materials and Methods for details). We then analyzed freshly isolated and processed samples directly from mice, digesting the resulting PDX tumoroids into a single cell suspension. Following the mass cytometry analysis, we generated a heatmap of the median surface expression of all the integrin subunits in our panel for each PDX model. Among these integrins, the surface expression of β1- and α2-integrin subunits (α2β1 is the main collagen I–binding integrin heterodimer) (Moreno-Layseca et al., 2019; Chastney et al., 2021) was markedly higher in the PDX#3 model compared to PDX#1 and PDX#2 (Fig. S2A). To visualize the overall integrin cell-surface profile, we combined a t-Distributed Stochastic Neighbor Embedding (tSNE; Amir et al., 2013) dimensionality reduction approach and a FlowSOM clustering (Quintelier et al., 2021). By doing so, it was possible to distinguish three populations depending on the expression of all the integrins in our panel (Fig. S2B). Overall, the PDX#3 model has a higher integrin expression level of all the considered subunits. By overlaying the β1-integrin and α2-integrin signal intensity to this clustering, we were able visualize the higher surface expression of β1-integrin and α2-integrin in PDX#3 (Fig. S2C). Next, we extracted the normalized surface expression values of β1- and α2-integrins for all the independent events. The histograms further demonstrated a marked variation in the cell surface integrin intensity and higher surface β1- and α2-integrin expression in the PDX#3 model compared to PDX#1 and PDX#2 (Fig. 2B).

To characterize the impact of the *ex vivo* culture on integrin surface expression, we performed a similar mass cytometry analysis, on tumoroids cultured three days in suspension or embedded in collagen for three days. These data recapitulated the integrin expression profiles of the cells directly extracted from the PDX tumors mentioned above (Fig. S2D). When plotting the corresponding histograms (Fig. 2C), we observed that collagen embedding of the PDX#3 tumoroids clearly increased the surface expression of β1- and α2-integrins in the cell population, in line with their ability to revert polarity in collagen (Fig. 2C). This prompted us to test the outcome of integrin inhibition on PDX#3 tumoroid polarity following embedding in collagen. Blocking both α2- and β1-integrins using established function-blocking antibodies (AIIB2 for β1-integrin and P1E6 for α2-integrin) (Yu et al., 2008; Berdichevsky et al., 1992) prevented the polarity reversion of PDX#3 in collagen (Fig. 2D-E). Furthermore, the outcome of integrin inhibition was evident in abolished tumoroid–collagen interactions and a round tumoroid morphology in live-imaging experiments (Fig. S2E-F, Video 2). Taken together, these data indicate that α2β1-integrin-mediated cell interaction with collagen-I and the ensuing downstream signaling are required for the phenotypic polarity reversion in the PDX#3 model (Fig. 2F).

### Polarity reversion is linked to altered expression of integrin traffic regulators

Whole exome sequencing revealed no mutations in ITGB1 and ITGA2 genes across the three PDXs (Fig. S2G), excluding a loss of protein function in PDX#1/PDX#2 as a potential explanation for the observed differences between the models. Cell-surface integrin levels are influenced not only by gene transcription but also by the dynamic process of integrin traffic, the continuous endocytosis of integrins from the plasma membrane and their return (recycling) (Caswell & Norman 2006; Bridgewater et al., 2012; Moreno-Layseca et al., 2019). To investigate the potential role of integrin trafficking in our models, we generated a list of 22 genes known to regulate this process and examined their expression in the same transcriptome data set (GSE152299) (Fig. 3A). PCA representation of this data (Fig. S3A) demonstrated a larger effect of matrix conditions on PDX#3 compared to PDX#1 and PDX#2. A Volcano plot representation of the gene set (Fig. 3B) identified two interesting targets for further investigation: *RAB11FIP1* and *SORL1*. *RAB11FIP1* encodes the Rab coupling protein (RCP; also called Rab11fip1, Rab11 family-interacting protein 1) which has been implicated in numerous studies as an important positive regulator of β1-integrin recycling, cancer cell invasion and metastasis (Caswell et al., 2008; Eva et al., 2010; Machesky, 2019). *SORL1* encodes the SorLA protein, a cell-surface sorting receptor implicated in trafficking of APP (Amyloid precursor protein) in neurons and rapid recycling of β1-integrins in breast cancer (Eggert et al., 2018; Pietilä et al., 2019). To validate the transcriptome data, we analyzed RCP and SorLA protein levels in the PDX tumoroids in suspension and collagen. RCP was downregulated upon collagen embedding on the protein level in all of the tumoroids and on the mRNA level in PDX#1 and PDX#3 (Fig. 3C-D). In contrast, *SORL1*/SorLA mRNA and protein expression were significantly upregulated only in collagen-embedded PDX#3 tumoroids (Fig. 3E-G and Fig. S3B). In breast cancer cells, integrin-ECM engagement activates the MAPK/ERK signaling pathway and ERK activity positively regulates *SORL1* transcription (Al-Akhrass et al., 2021; Al-Akhrass et al., 2022), which could also be a possible mechanism for collagen-induced upregulation of *SORL1*/SorLA in mucinous CRC. Indeed, we detected significantly elevated ERK phosphorylation in PDX#3 tumoroids in collagen (Fig. 3H and S3C) and collagen-induced SorLA upregulation was sensitive to integrin function-blocking antibodies (Fig. 3I and S3D). These data suggest the possibility that the PDX#3 tumoroids in collagen switch from RCP-mediated long-loop integrin recycling to a SorLA-driven rapid recycling pathway in a manner that is dependent on initial matrix-induced integrin signaling.

**FIG3.**
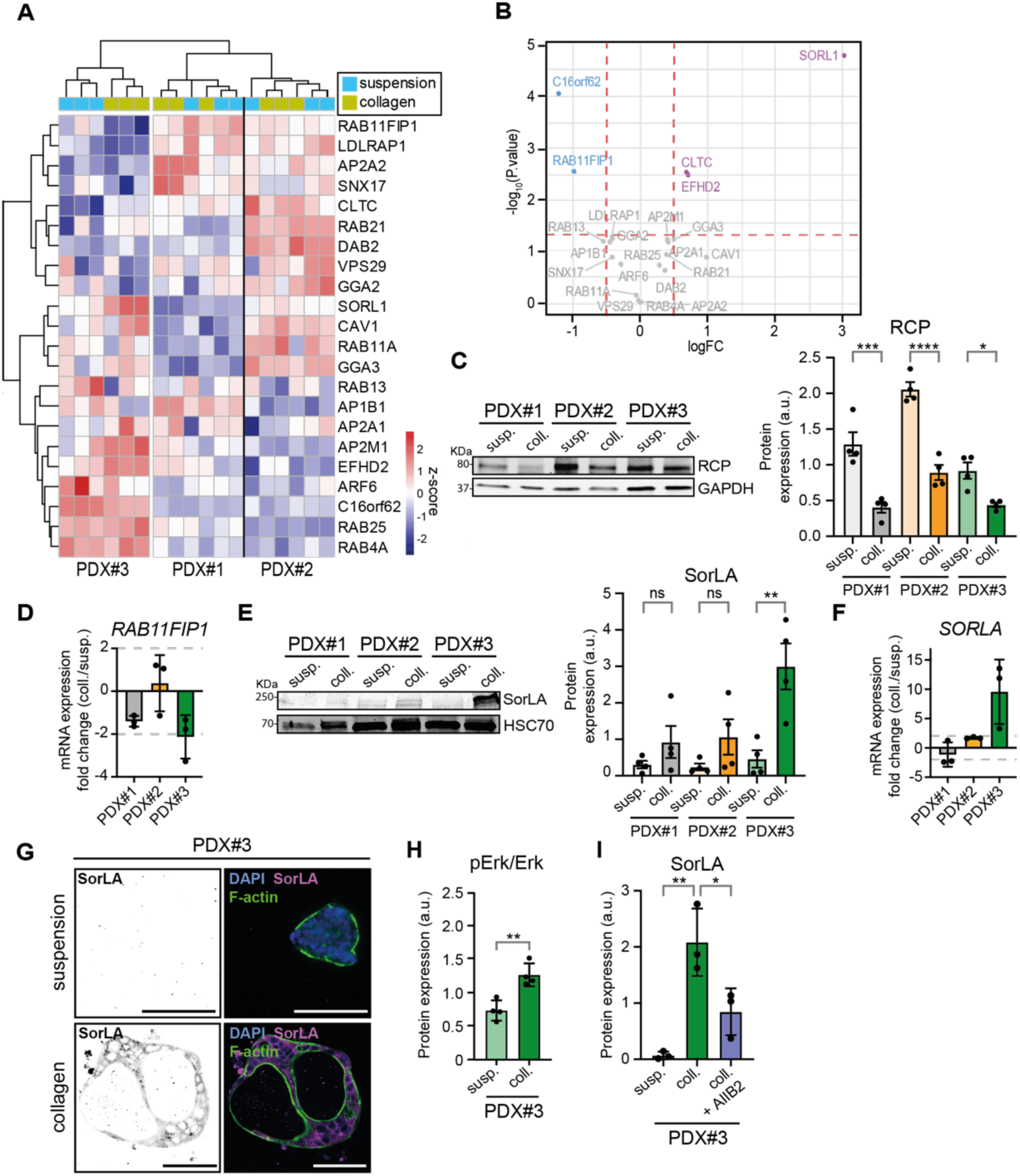
Key integrin-β1 trafficking regulators are differentially expressed among PDX models and matrix conditions. (A) Relative expression of key regulators in integrin-β1 trafficking, in PDX#1-, PDX#2- and PDX#3-generated tumoroids in suspension and in collagen. (B) Volcano plot generated for the same geneset comparing PDX#3-generated tumoroids in collagen vs. suspension. Significantly upregulated genes in collagen are in purple, downregulated genes are in blue. (C) Western-blot analysis of RCP and GAPDH (loading control) and corresponding quantifications of RCP in PDX#1-, PDX#2- and PDX#3-generated tumoroids in suspension and in collagen, normalized to the loading control (n=4, One-way ANOVA with Tukey’s multiple comparison test, p-value_PDX#1 coll vs. susp_=0.0002, p-value_PDX#2coll vs. susp_<0.0001, p-value_PDX#3 coll vs. susp_=0.0492). (D) Change in *RAB11FIP1* (gene encoding for RCP) mRNA expression in collagen vs. suspension from mRNA microarray analysis of PDX#1-, PDX#2- and PDX#3-generated tumoroids. (E) Western-blot analysis of SorLA and HSC70 (loading control) and corresponding quantifications of SorLA in PDX#1-, PDX#2- and PDX#3-generated tumoroids in suspension and in collagen normalized to loading control (n=4, One-way ANOVA with Tukey’s multiple comparison test, p-value_PDX#1 coll vs. susp_=0.8651, p-value_PDX#2 coll vs. susp_=0.6725, p-value_PDX#3 coll vs. susp_=0.0024). (F) Change in SORL1 mRNA expression in collagen vs. suspension based on mRNA microarray analysis of PDX#1-, PDX#2- and PDX#3-generated tumoroids. (G) Immunofluorescence of PDX#3-generated tumoroids in suspension and in collagen, showing SorLA. (H) Western-blot quantifications of Erk phosphorylation ratio in PDX#3 in suspension and collagen (n=4, paired t-test, p-value=0.0036). (I) Western-blot quantifications of SorLA expression in PDX#3 in suspension, collagen, and in collagen while treated with AIIB2 (n=3, One-way ANOVA with Tukey’s multiple comparison test, p-value_coll. vs. susp.=_0.0027, p-value_coll.+AIIB2 vs. coll.=_0.0271). Scale bars, 100 μm.

### Apical-in polarity in collagen is established through SorLA-dependent β1-integrin recycling

To explore SorLA-mediated β1-integrin regulation, we first investigated the impact of collagen-embedding on β1-integrin mRNA and total protein levels (Fig. 4A-B). Western-blot analysis of β1-integrin typically reveals two bands, a slower-migrating band representing the mature, fully-glycosylated protein, and a faster-migrating band, indicating the immature, ER-resident protein. The relative abundance of these two protein forms has been linked to alterations in integrin traffic, with a higher ratio of the slower-migrating band corresponding to increased β1-integrin recycling and plasma membrane residency time (Böttcher et al., 2012). Interestingly, overall β1-integrin levels were not affected in any of the three PDX tumoroids in response to collagen-I. However, quantification of the ratio of the mature versus immature protein form revealed a significant increase in integrin maturation in the PDX#3 model in response to collagen-I. The identity of the slower migrating band as mature β1-integrin was further confirmed with digestion of the lysates with PNGase (Fig. S4A).

**FIG4.**
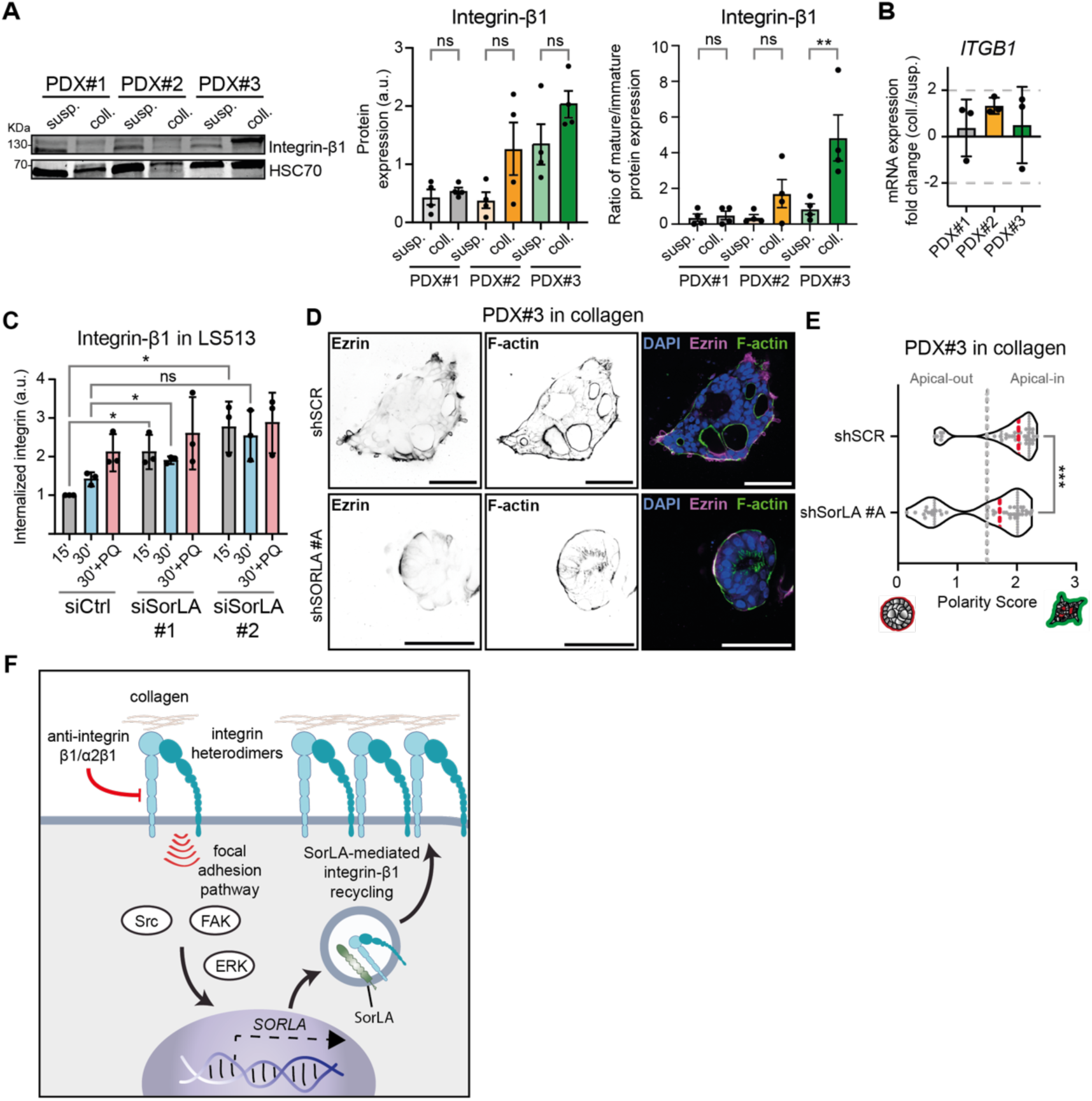
SorLA-mediated integrin-β1 trafficking is necessary for polarity reversion in collagen. (A) Western-blot and corresponding quantifications and maturation ratios of integrin-β1 in PDX#1-, PDX#2- and PDX#3-generated tumoroids in suspension and in collagen (for total expression: n=4, One-way ANOVA with Tukey’s multiple comparison test, p-value_PDX#1 coll vs. susp_=0.9996, p-value_PDX#2 coll vs. susp_=0.2173, p-value_PDX#3 coll vs. susp_=0.4623; for maturation ratio: n=4, One-way ANOVA with Tukey’s multiple comparison test, p-value_PDX#1 coll vs. susp_>0.9999, p-value_PDX#2 coll vs. susp_=0.6903, p-value_PDX#3 coll vs. susp_=0.0043). (B) ITGB1 mRNA expression fold change in collagen vs. suspension generated from an RNA microarray analysis of PDX#1-, PDX#2- and PDX#3-generated tumoroids. (C) Integrin-β1 uptake assay in LS513 after siRNA-induced silencing of SorLA. Timepoints are 15’ and 30’, and the recycling inhibitor used is primaquine (PQ) (n=3, independent Welch’s t-tests, p-value_15’ si#1 vs. siCtrl_=0.0432, p-value_15’ si#2 vs. siCtrl_=0.0406, p-value_30’ si#1 vs. siCtrl_=0.0172, p-value_30’ si#2 vs. siCtrl_=0.0877). (D) Immunofluorescence of PDX#1-, PDX#2- and PDX#3-generated tumoroids in suspension and in collagen, showing their polarity status through Ezrin staining after KD of SorLA by LV shRNA. (E) Comparison of polarity scores of PDX#3 in collagen after SorLA silencing. (N_shSCR_=38, N_shSORLA #A_=44, Mann-Whitney test, p-value_shSORLA #A vs. shSCR_=0.0007). (F) Proposed mechanism of collagen-induced Erk, FAK and Src activation and SorLA expression, leading to integrin-β1 recycling. Scale bars, 100 μm.

To formally assess the role of SorLA in integrin recycling within MUC CRC cells, we sought to establish a more tractable cell-based model, given the technical challenges associated with performing cell biology with PDX models. Similarly to some other cell lines (Lubarsky et al., 2003), the MUC CRC cell line LS513 generates buds from confluent cultures and secretes TSIPs in the medium. We observed that these cells phenocopied all the key features of the PDX#3 tumoroids. Once embedded in collagen, the LS513-generated TSIPs formed lumens, shifted significantly towards an apical-in polarity (Fig. S4B-C) and upregulated SorLA levels (Fig. S4D). We concluded that these cells are a good surrogate to study β1-integrin traffic. Previous work has shown that β1-integrins recycle to the plasma membrane with a turnover rate of 10–15 min (Argenzio et al., 2014; Diggins et al., 2018; Dozynkiewicz et al., 2012). Impaired recycling can lead to the gradual intracellular accumulation of β1-integrin (Sahgal et al., 2019). We silenced *SORL1* in LS513 with two independent siRNA oligos (Fig. S4E) and used the gold-standard cell-surface biotinylation-based integrin uptake assay to monitor endocytosed β1-integrin levels (Arjonen et al., 2012; Farage et al., 2021). *SORL1* silencing significantly elevated intracellular β1-integrin levels within 15 minutes, mirroring the effects of the recycling inhibitor primaquine (Fig. 4C). This data indicates that loss of SorLA, similar to our previous findings in breast cancer (Pietilä et al., 2019), primarily increases intracellular β1-integrin levels by inhibition of recycling.

To functionally test the role of SorLA in collagen-induced polarity reversion, we established a patient-derived organoid line (PDO line) as described in Cartry et al., 2023 from the PDX#3 tumor (PDX#3 PDO line) and silenced *SORL1* using lentiviral shRNA. We obtained more than 50% silencing (Fig. S4F) and observed a significant reduction in PDX#3 polarity reversion in collagen-I, with clusters retaining a mostly apical-out polarity (Fig. 4D-E).

Taken together, the data thus far are supportive of a positive feedback loop (Fig. 4F) whereby α2β1-integrin adhesion initiates signaling from collagen through Src, FAK and ERK, upregulating SorLA expression. This induces rapid integrin recycling, giving rise to higher integrin cell-surface localization that supports enhanced cell–collagen interaction and apical-in polarity.

### SorLA-dependent Integrin-β1 trafficking is induced by HER2/HER3 signaling

The human epidermal growth factor receptor 2 (HER2) is a transmembrane receptor tyrosine kinase (RTK) implicated in breast and gastroesophageal cancers, where targeted therapies against this RTK have already been developed for clinical use (Bartley et al., 2016). HER2 is also an emerging biomarker in CRC (Ivanova et al., 2022). We have shown that SorLA interacts with HER2 homo- and HER2/HER3 heterodimers supporting rapid HER2/HER3 recycling and protein stability and oncogenic signaling in breast and bladder cancers (Pietilä et al., 2019; Al-Akhrass et al., 2021). Furthermore, Heregulin-β1-induced HER2/HER3 signaling to ERK increases SorLA transcription (Al-Akhrass et al., 2021). To explore whether RTK signaling is implicated in MUC CRC polarity, we analyzed HER2 and HER3 expression in PDX#1, PDX#2 and PDX#3 tumoroids in suspension and collagen. Although HER2 is upregulated in both PDX#2 and PDX#3, only the collagen-responsive PDX#3 clearly upregulated both HER2 and HER3 protein expression simultaneously when embedded in collagen (Fig. 5A-B). Immunofluorescence staining further validated these results with clear simultaneous HER2 and HER3 signal detected only in PDX#3 in collagen, where HER2 mostly localized at the basal pole (Fig. 5C and Fig. S5A). Similar to the collagen-induced SorLA expression, HER2 and HER3 upregulation were also integrin dependent in PDX#3 in collagen (Fig. 5D-E and Fig. S5B).

**FIG5.**
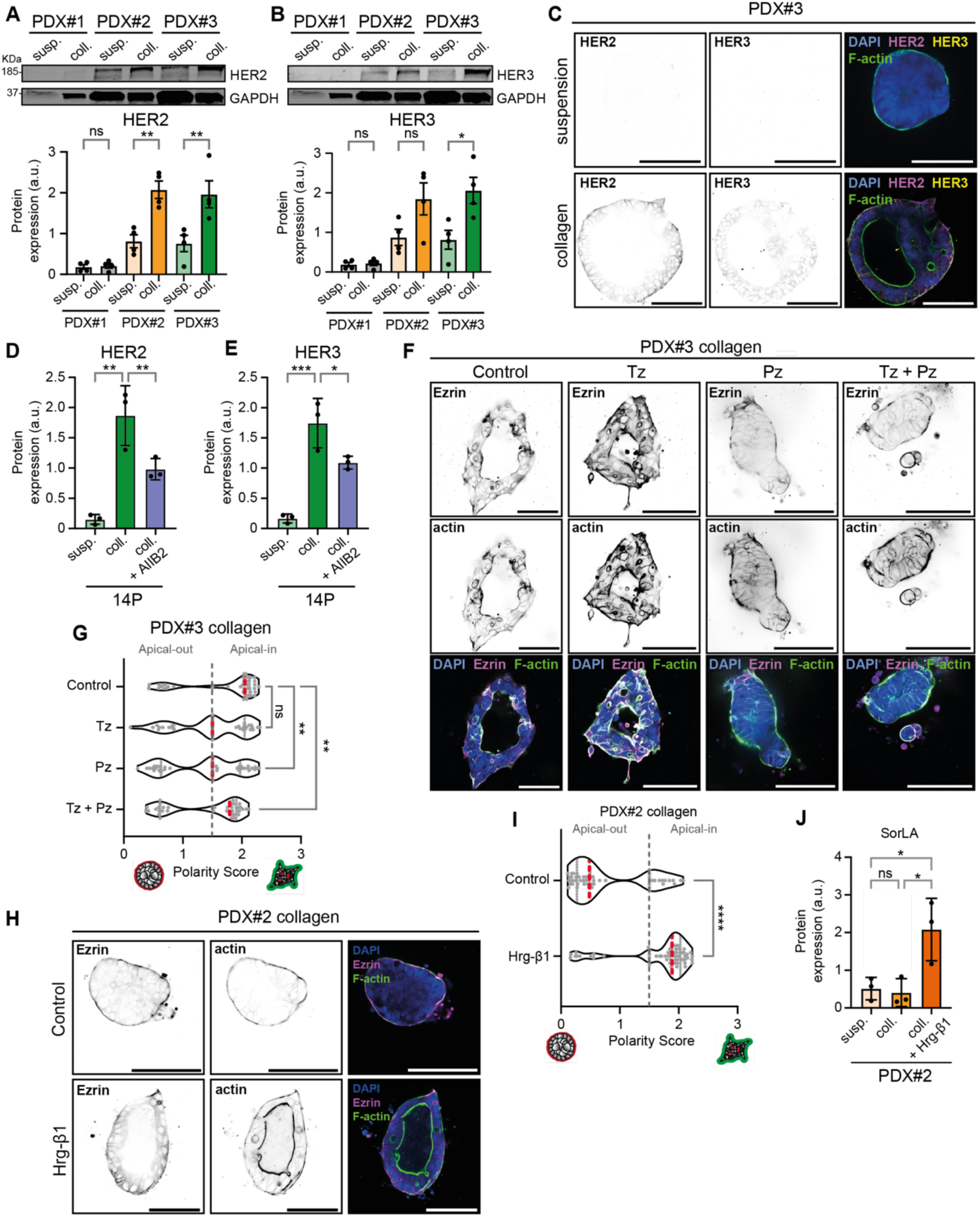
HER2 and HER3 control apicobasal polarity orientation. (A) Western-blot and corresponding quantifications of HER2 in PDX#1-, PDX#2- and PDX#3-generated tumoroids in suspension and in collagen (for total expression: n=4, One-way ANOVA with Tukey’s multiple comparison test, p-value_PDX#1 coll vs. susp_>0.9999, p-value_PDX#2 coll vs. susp_=0.0027, p-value_PDX#3 coll vs. susp_=0.0042). (B) Western-blot and corresponding quantifications of HER3 in PDX#1-, PDX#2- and PDX#3-generated tumoroids in suspension and in collagen (for total expression: n=4, One-way ANOVA with Tukey’s multiple comparison test, p-value_PDX#1 coll vs. susp_>0.9999, p-value_PDX#2 coll vs. susp_=0.1165, p-value_PDX#3 coll vs. susp_=0.0264). HER2 and HER3 have been blotted on the same membrane and were therefore normalized to the same GAPDH signal. (C) Immunofluorescence of PDX#3-generated tumoroids in suspension and in collagen, showing HER2 and HER3. (D) Western-blot quantifications of HER2 expression in PDX#3 in suspension, collagen, and collagen treated with AIIB2 (n=3, One-way ANOVA with Tukey’s multiple comparison test, p-value_coll. vs. susp.=_0.0011, p-value_coll.+AIIB2 vs. coll.=_0.0282). (E) Western-blot quantifications of HER3 expression in PDX#3 in suspension, collagen, and collagen treated with AIIB2 (n=3, One-way ANOVA with Tukey’s multiple comparison tests, p-value_coll. vs. susp.=_0.0006, p-value_coll.+AIIB2 vs. coll.=_0.0403). (F) Immunofluorescence of PDX#3-generated tumoroids in suspension, collagen, and in collagen while treated with a HER2/HER2 homodimerization blocking antibody (Trastuzumab, Tz, 10 μg/ml), a HER2/HER3 heterodimerization blocking antibody (Pertuzumab, Pz, 10 μg/ml) or a combination of both (Tz+Pz). Polarity status is depicted though Ezrin staining. (G) Comparison of polarity scores in PDX#3 in collagen with HER2/HER3-blocking antibodies (n_Ctrl_=44, n_Tz_=39, n_Pz_=47, n_Tz+Pz_=38, One-way ANOVA, p-value_Tz vs. Ctrl_=0.2086, p-value_Pz vs. Ctrl_=0.0063, p-value_Pz+Tz vs. Ctrl_=0.0042). (H) Immunofluorescence of PDX#2-generated tumoroids in suspension, collagen, and in collagen while treated with a HER3 ligand (Heregulin-β1, Hrg-β1, 20 ng/ml), showing their polarity status through Ezrin staining. (I) Comparison of polarity scores in PDX#2 in collagen with, treated with Hrg-β1 (n_Ctrl_=56, n_Hrg-β1_=62, Mann-Whitney test, p-value<0.0001). (J) Western-blot quantifications of SorLA expression in PDX#2 in suspension, collagen, and in collagen while treated with Hrg-β1 (n=3, One-way ANOVA with Tukey’s multiple comparison test, p-value_susp. vs. coll+ Hrg-β1 =_0.0305, p-value_coll.+Hrg-β1 vs. coll._=0.0231). Scale bars, 100 μm.

To assess the impact of HER2 and HER3 signaling on tumoroid polarity, we used two monoclonal antibodies: trastuzumab (prevents HER2 homodimerization and autophosphorylation) and pertuzumab (prevents HER2 and HER3 heterodimerization), both of which are in clinical use for breast cancer treatment (Swain et al., 2015). In our set-up, trastuzumab alone had no significant effect on PDX#3 polarity. However, pertuzumab alone or in combination with trastuzumab significantly shifted PDX#3 tumoroids toward an apical-out polarity (Fig. 5F-G), indicating a functional role for HER2/HER3 signaling, but not HER2/HER2 homodimers, in PDX#3 polarity reversion in response to collagen. Conversely, HER2/HER3 activation with the HER3 ligand Heregulin-β1 was sufficient to trigger collagen-induced polarity reversion in the thus far matrix agnostic PDX#2 tumoroids (Fig. 5H-I). Upon HER3 stimulation with Heregulin-β1, SorLA was also found to be upregulated in PDX#2 (Fig. 5J and S5E). The PDX#1 tumoroids did not respond, presumably due to their lack of HER3 expression (Fig. 5B and S5C-D). Additionally, whole exome sequencing (Fig. S5F) indicated that there are no mutations in the aforementioned genes, *RAB11FIP1*, *SORL1*, *HER2* and *HER3*, except for a missense mutation in HER2 in PDX#2 (a G>A at nucleotide 1943 encoding Arg648Gln), which SIFT and PolyPhen scores suggest is benign on its structure and function (Flanagan et al., 2010; Adzhubei et al., 2013).

### Apical-in CRC polarity is associated with enhanced collagen remodeling

Since the reversion of PDX#3 tumoroid polarity in collagen was dependent on direct integrin-mediated cell-ECM interactions, we wanted to investigate whether this would also lead to concomitant remodeling of the ECM itself. To study the mechanics of the SorLA/HER2/HER3-dependent polarity reversion, we visualized collagen fiber rearrangements using a fluorescent probe specific for fibrillar collagen (mScarlet-conjugated CNA35) (Aper et al., 2014). The PDX#3 tumoroids remodeled the collagen matrix resulting in increased fiber alignment in the tumoroid-proximal ECM (Fig. 6A-B and Fig. S6A), whereas integrin inhibition with AIIB2 antibody or dual inhibition of HER2/HER3, with the combination of pertuzumab and trastuzumab, significantly decreased matrix alignment (Fig. 6A-B and Fig. S6A). Conversely, treating PDX#2 tumoroids with Heregulin-β1 increased collagen fiber orientation, indicating elevated collagen remodeling (Fig. 6A-B and Fig. S6B).

**FIG6.**
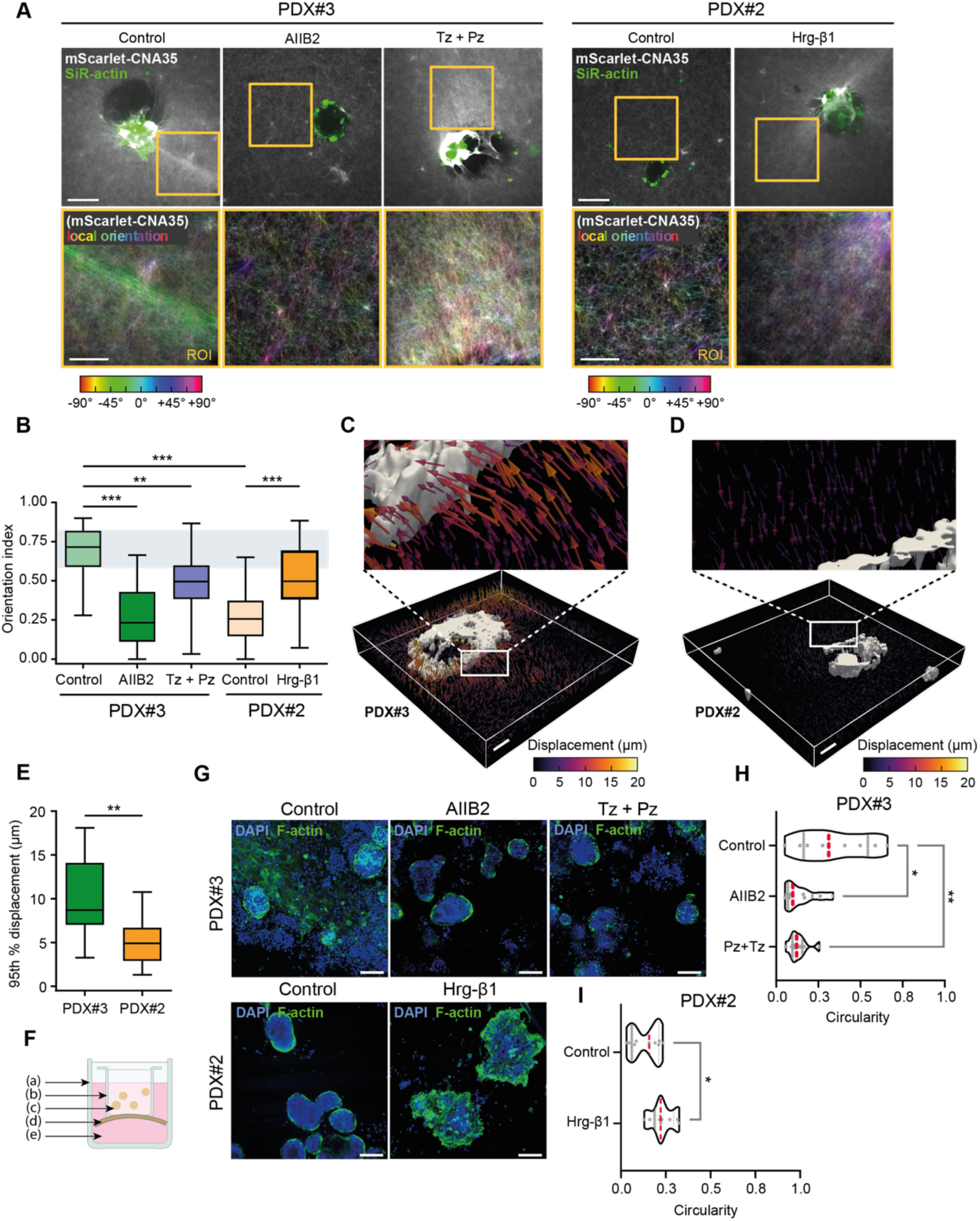
Collagen remodeling is correlated to apicobasal polarity status in collagen. (A) Visualization of collagen fiber orientations proximal to PDX#2 and PDX#3 tumoroids treated with integrin-β1-blocking antibody (AIIB2), HER2/HER3-blocking antibodies (Pz+Tz) and HER3 ligand (Hrg-β1). Collagen fibers are marked with mScarlet-CNA35, and the images are color-coded based on the local orientation of the fibers. Scale bars, 50 μm (main), 20 μm (inset) (B) Collagen fiber orientation index in PDX#2 and PDX#3 treated as indicated (n_PDX#3 Ctrl_=28, n_PDX#3 AIIB2_=28, n_PDX#3 Pz+Tz_=21, n_PDX#2 Ctrl_=29, n_Hrg-β1_=23, One way ANOVA with Šidák’s multiple comparison test, p-value_PDX#3AIIB2 vs. Ctrl_<0.0001, p-valuePDX#3_Pz+Tz vs. Ctrl_=0.0008, p-value PDX#2_Hrg-β1 vs. Ctrl_<0.0001, p-value_PDX#3 vs. PDX#2_<0.0001). (C) Visualization of reversible displacements exerted on the collagen matrix by a PDX#3 tumoroid (rendered in the imaged using thresholded F-actin signal). (D) Visualization of reversible displacements exerted on the collagen matrix by a PDX#2 tumoroid. (E) Quantification of the matrix displacements exerted by the PDX#3 and PDX#2 tumoroids (n_PDX#3_=27, n_PDX#2_=29, Mann-Whitney test, p-value=0.0032). (F) Set-up for the peritoneum invasion assay: (a): well of a 12-well cell culture plate, (b) Transwell cell invasion insert, from which the membrane has been removed, (c) tumoroids, (d) decellularized peritoneum, (e) medium. (G) Fluorescence imaging of PDX#2- and PDX#3-generated tumoroids after 6 days of cell contact with the decellularized peritoneum, after treatment with AIIB2 (1 μg/ml), Pertuzumab+Trastuzumab (Tz+Pz, 10 μg/ml, 10 μg/ml) or Hrg-β1 (20 ng/ml). (H) Comparison of polarity scores in PDX#3 in collagen after treatment with AIIB2 (1 μg/ml) or Pertuzumab+Trastuzumab (Tz+Pz, 10 μg/ml, 10 μg/ml) on 9 spheres selected randomly (Mann-Whitney test, p_Ctrl. Vs. Pz+Tz_=0.0073, p_Ctrl. Vs. AIIB2_=0.0121). (I) Comparison of circularity in PDX#2 in collagen after treatment with Heregulin-β1 (Hrg-β1, 20 ng/ml) on 9 spheres selected randomly (Mann-Whitney test, p-value_Hrg-β1 vs. Ctrl_=0.0142). Scale bars, 100 μm (main), 50 μm (3D projections). Vector scaling is set to 200% for clarity.

To further investigate biophysical interaction between the cells and their ECM, we imaged collagen-embedded PDX#3 and PDX#2 tumoroids before and after Latrunculin B treatment. Latrunculin B disrupts the actin cytoskeleton, releasing cellular tension and leading to a recoil in the tensed matrix (Fig. 6C-D and Fig. S6C). Comparing the images before and after treatment allowed us to compute the reversible displacement of collagen fibers caused by tumoroid contraction in each condition. Using a previously described MATLAB pipeline (Barrasa-Fano et al., 2021a), we observed significantly higher reversible collagen displacements proximal to the PDX#3 tumoroids compared to PDX#2 (Fig. 6C-E). This suggests

that the collagen fibers are under constant mechanical tension exerted by the MUC CRC cells, and such forces are much higher in the apical-in tumoroids.

### HER2/HER3 and integrins regulate CRC tumoroid interaction with the peritoneum *ex vivo*

CRC often metastasizes to the liver and peritoneum and successful metastasis depends on the cancer cells’ ability to adhere to and invade this tissue (Barbazán et al., 2017). To study peritoneal invasion with high granularity and in response to various treatments, we set up an in vitro model that recapitulates CRC clusters adhering to and invading the surface of the peritoneum. We isolated and de-cellularized mouse peritoneum, a technique previously applied to the mouse mesentery (Glentis et al., 2017), which was shown to recapitulate the organization and characteristics of the in vivo basement membrane (Varinelli et al., 2022). We plated the PDX tumoroids on the de-cellularized peritoneum (Fig. 6F) and tested the outcome of different treatments (AIIB2 or Trastuzumab+Pertuzumab for PDX#3, Heregulin-β1 for PDX#2) on the ability of the tumoroids to adhere and spread on the tissue. Despite the relatively low adhesion rate, we observed tumor spheres adhering to the peritoneum (Fig. 6G) and based on actin staining quantified their circularity. The PDX#3 tumoroids, which revert their polarity to apical-in when embedded in collagen, spread extensively on the peritoneum with individual cells dissociating from the clusters. Integrin- and HER2/HER3-inhibiting antibodies significantly blocked this (Fig. 6G-H). In contrast, the untreated PDX#2 tumoroids did not display extensive spreading on the peritoneum. However, stimulation with the HER3 ligand Heregulin-β1 enhanced spreading and reduced spheroid circularity (Fig. 6G-I).

Taken together, these data are concordant with a model where HER2 and HER3 regulate integrin-collagen interactions in MUC CRC tumoroids, and increased integrin cell surface availability and signaling are associated with both apical-in polarity and altered tumoroid-proximal ECM architecture.

### High HER2/HER3/SORLA expression correlates with apical-in polarity of CRC tumors

To determine if the *ex vivo* findings on SorLA, HER2, and HER3 were applicable *in vivo*, we investigated their correlation in PDX tumors in mice. We observed high HER2 staining and predominantly apical-in polarity in the PDX#3 tumors. Conversely, the staining intensity was low in the predominantly apical-out PDX#2 tumors (Fig. S7A). Encouraged by the concordant results between *in vivo* and *ex vivo* models regarding HER2, we stained a cohort of 25 human MUC CRC patient samples (Barresi et al., 2015) for these three proteins. The staining revealed a strong positive signal for HER2/SorLA and HER2/HER3 in the normal epithelium as well as the epithelium of apical-in CRC clusters whereas in apical-out structures the staining intensities of SorLA and HER2 were lower (Fig. 7A and Fig. S7B). The quantifications revealed interesting findings: first, there is a positive correlation between HER2 and HER3, as well as between HER2 and SorLA, both in apical-in and apical-out structures (Fig. 7B). Second, there is a modest correlation between the polarity status and HER2 expression (higher in apical-in than in apical-out). Although similar tendencies were seen for HER3 and SorLA, the lack of statistical significance did not allow us to draw any strong conclusions (Fig. 7C).

**FIG7.**
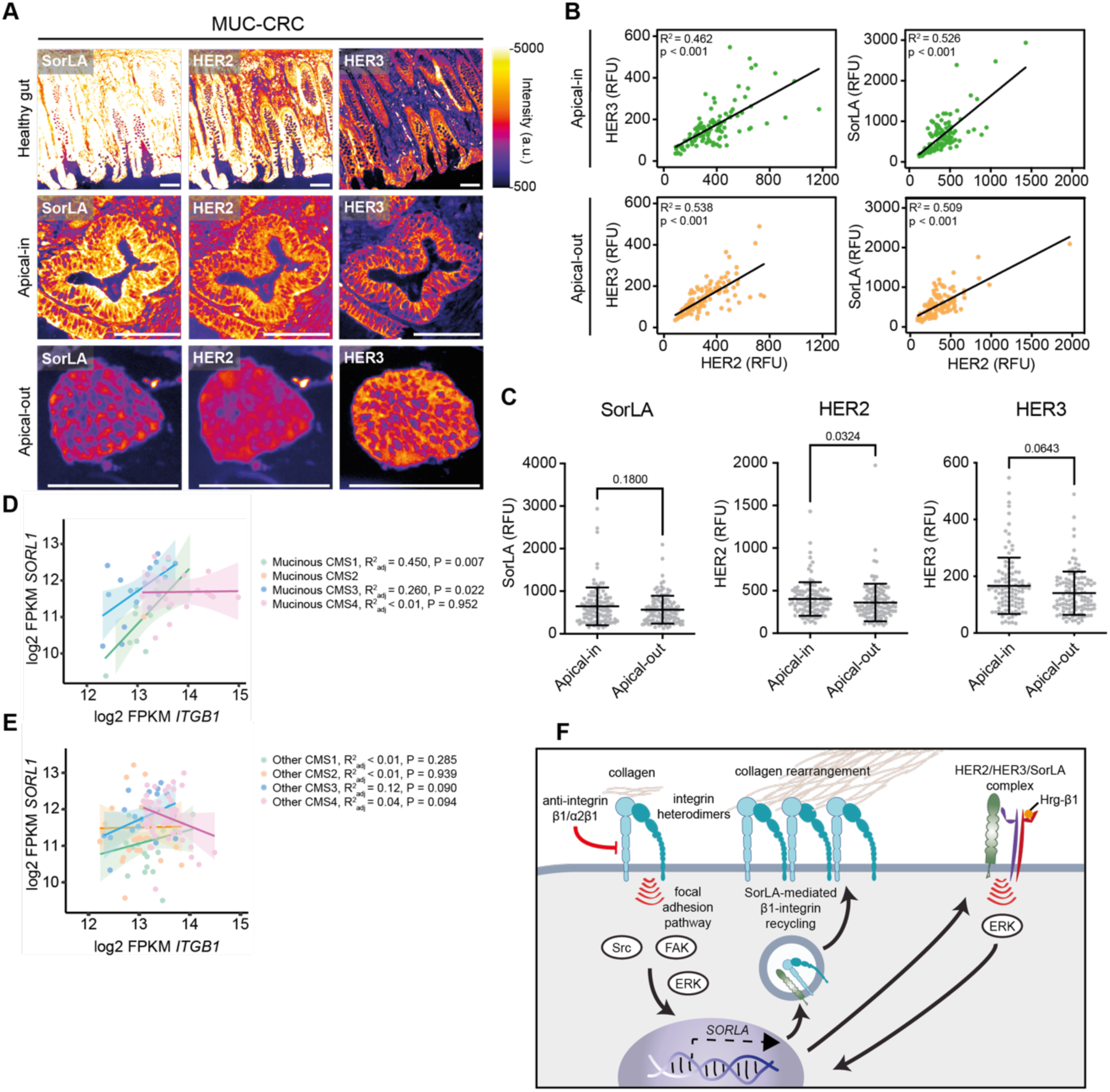
HER2, HER3 and SorLA expressions are correlated *in vivo*. (A) Immunohistochemistry of healthy gut tissue, apical-in and apical-out cancer structures from MUC CRC patients stained for HER2, HER3 and SorLA. These panels have been made from two consecutive different slides that were cut at different depths of the tumor, which explains the slight change in structure between the pictures. (B-C) Correlation plots (from a simple linear regression) (B) and expression (C) for HER2, HER3 and SorLA in a cohort of 25 patients (5 apical-in and 5 apical out structures chosen for each patient when possible) in apical-in and apical-out cancer structures. (HER2: N_Apical-in_=115, N_Apical-out_=120, Mann-Whitney test; HER3: N_Apical-in_=110, N_Apical-out_=120, Mann-Whitney test; SorLA: N_Apical-in_=115, N_Apical-out_=120, Mann-Whitney test). (D) Correlation (from a simple linear regression) between *ITGB1* and *SORL1* expression in mucinous CRCs of CMS1-4 (log2 of Fragments per Kilobase of transcript per Million mapped reads (FPKM) plotted), (E) Correlation between *ITGB1* and *SORL1* expression in non-mucinous CRCs of CMS1-4 (log2 of Fragments per Kilobase of transcript per Million mapped reads (FPKM) plotted), (F) Proposed mechanism of collagen-induced Erk, FAK and Src activation and SorLA expression, leading to integrin-β1 recycling. This mechanism is supported by a HER2/HER3/SorLA positive feedback loop and results in efficient matrix-sensing and collagen rearrangement. Scale bars, 100 μm.

Interestingly, by analyzing gene expression profiles from a TCGA CRC cohort which histological characterization has previously been performed (Nguyen et al., 2021), and dividing them into two groups (mucinous and non-mucinous) (Fig. S7C), we were able to see a positive correlation between *ITGB1* and *SORL1* mRNA expression both in CMS1 and CMS3 (Consensus Molecular Subtype 1 and 3) solely for MUC CRCs (CMS2 being mostly non-represented in the mucinous CRCs), which was not the case for the non-mucinous CRCs (Fig. 7D-E).

Cumulatively, using cell- and mechanobiology methods to interrogate MUC CRC PDX tumors ex-vivo and patient samples we report a novel polarity determination mechanism whereby HER2/HER3/SorLA-controlled −β1-integrin traffic controls tumor-ECM interactions, ECM rearrangements and spreading onto the peritoneum (Fig. 7F).

## Discussion

Epithelial polarity is often considered a feature of normal tissue, maintained by cell-cell and cell-basement membrane interactions, and which is progressively lost in cancer in conjunction with increased invasion and metastasis. Our study uncovers a dynamic crosstalk of integrins, HER-family receptor tyrosine kinases and receptor membrane traffic orchestrated by SorLA that jointly determine MUC CRC polarity. Using cell- and mechanobiology methods to interrogate *ex vivo* CRC PDX tumors and patient samples, we report a novel polarity determination mechanism whereby HER2/HER3/SorLA-controlled β1-integrin traffic regulates tumor-ECM interactions, ECM remodeling and cancer cell dissemination on the peritoneum. Furthermore, expression of these receptors shows significant positive correlation in clinical patient specimens (Fig 7B).

SorLA is a well-established sorting protein regulating membrane traffic of APP in neurons, insulin receptor in adipocytes and HER2/HER3 receptors specifically in HER2-dependent breast cancer (Al-Akhrass et al., 2021; Pietilä et al., 2019; Andersen et al., 2005, 2006; Klinger et al., 2011; Spoelgen et al., 2009; Schmidt et al., 2016; Whittle et al., 2015). To the best of our knowledge, SorLA has not been previously implicated in CRC or in the regulation of cell polarity. We find that SorLA is a central nexus for polarity determination in MUC CRC TSIPs. SorLA levels are low in TSIPs is suspension but become rapidly upregulated in integrin-high TSIPs capable of forming initial contacts with collagen and adopting apical-in topology. ECM contact triggers two regulatory feedback loops: enhanced integrin recycling supportive of polarity maintenance and increased HER2/HER3 protein levels that promote SorLA expression. Conversely, inhibition of integrins or HER2/HER3 receptors or silencing of SorLA impairs ECM-induced polarity reversion of TSIPs. These data increase our understanding of MUC CRC cluster behavior during cancer progression and are likely to have important functional implications, as previous work has indicated that non-reverted CRC clusters are capable of long-distance collective ameboid migration and are more resistant to chemotherapy (Pagès et al., 2022; Ashley et al., 2019). On the other hand, we observed that efficient polarity reversion and apical-in morphology facilitated MUC CRC interaction with decellularized mouse peritoneum and promoted local collagen remodeling. Similar radially organized ECM signature drives local invasion and is associated with poor patient outcomes in breast cancer (Provenzano et al., 2006; Conklin et al., 2011), suggesting that the impact of polarity reversion in MUC CRC may be multifaceted and its role during the different phases of the metastatic cascade and in different tissue microenvironments warrants careful investigation.

In normal and cancer cells, integrins are constantly trafficked from the plasma membrane to endosomes and recycled back to facilitate dynamic cell adhesion (Moreno-Layseca et al., 2019; Paul et al., 2015). Increased integrin traffic facilitates invasion and metastasis in many cancer types, including breast and pancreatic cancer (Caswell et al., 2006; De Franceschi et al., 2015). In particular, increased integrin recycling via RCP has been linked to β1-integrin recycling in complex with EGFR to drive cancer cell motility in 3D matrix (Machesky, 2019; Caswell et al., 2008; Rainero et al., 2012; Jacquemet et al., 2013; Muller et al., 2009). Here we find that MUC CRC tumor spheres, interacting with collagen, significantly downregulate RCP expression and upregulate SorLA. This implies that in reverted polarity TSIPs’ integrin recycling is switched to a polarity-maintaining receptor traffic program supportive of HER2/HER3 signaling.

Integrin expression is frequently altered in CRC, with laminin-binding α6β4-integrin being frequently overexpressed and perhaps the most broadly studied integrin heterodimer (Beaulieu et al., 2020). Of the collagen-binding integrins, α1β1-integrin is normally restricted to the crypt epithelium but is upregulated in 65% of CRCs where its expression is driven by MYC (Boudjadi et al., 2013, 2016). Surprisingly, the main fibrillar-collagen binding integrin-α2β1 has not been investigated extensively in CRC. We find here that both α1β1- and α2β1-integrins are expressed in the PDX-tumors and tumor spheres, with α2β1 having the highest expression in the apical-in polarized PDX model. Inhibitory antibodies for α2-integrin are sufficient to block collagen-induced polarity reversion, implying that α1β1-collagen interaction alone is not supportive of polarity reversion. This is an interesting distinction which could be linked to the pro-tumorigenic role of α1β1-integrin in CRC and warrants further investigation in the future.

Anti-EGFR therapy is in clinical use for the treatment of CRC, whereas trials for targeted therapies against HER2 have not resulted in approval of anti-HER2 therapy in CRC (Ye et al., 2022; Nowak et al., 2020). Interestingly, HER2 amplifications and elevated levels of the HER3 ligand, Heregulin-β1, are linked to resistance of CRC to anti-EGFR therapeutic antibody cetuximab (Martin et al., 2013; Yonesaka et al., 2011). As expected, distinct CRC subtypes show differential responses to cancer therapeutics owing to their different biology. Here, we find a correlation between *ITGB1* and *SORL1* expressions solely in the mucinous subtypes of CRC, which shows the presented mechanism might be limited to specific histological phenotypes of cancer. Our discovery outlining the polarity regulation of MUC CRC by HER2/HER3/SorLA signaling *ex vivo* and in human tumors suggests a functionally important role for the pathway in this CRC subtype, and may warrant further attention in the field of CRC therapy.

## Acknowledgements

We thank P. Laasola and J. Siivonen for technical assistance and the Ivaska lab and Jaulin lab for scientific discussions. The Cell Imaging and Cytometry Core (Turku Bioscience, University of Turku, Åbo Akademi University and Biocenter Finland), Turku Bioimaging and Turku Center for Disease Modelling (TCDM) are acknowledged for services, instrumentation and expertise. This work was supported by the Finnish Cancer Institute (K. Albin Johansson Professorship to J.I.); a Research Council of Finland research project (grant no. 325464 to J.I.) and Centre of Excellence program (grant no. 346131 to J.I.); the Cancer Foundation Finland (to J.I.); the Sigrid Juselius Foundation (to J.I.); and the Research Council of Finland InFLAMES Flagship program (grant no. 337530 and 357910). This work received support from the Institut National du Cancer (INCa) (grant 2020-1-PLBIO-04-IGR-1), fundraising against CRC and Mars Bleu from the Gustave Roussy foundation. This work has also benefited from a government grant managed by the National Research Agency under the 5^th^ PIA, integrated into France 2030 with the reference ANR-21-RHUS-0003. N.P. was supported by the “PhD in Oncology” program of the Fondation Philantropia, as well as the Pentti and Tyyni Ekbom Fund. A.I. was supported by the Finnish Cultural Foundation, the Orion Research Foundation and the K. Albin Johansson’s Foundation. G.F. was supported by the Research Council of Finland (grant 332402) and TCSMT postdoctoral research grants. I.B., J.S. and P.K. (from Turku Bioimaging, part of the Finnish Advanced Microscopy Node of Euro-Bioimaging Finland) were funded by the Research Council of Finland, FIRI 2023 (grant no. 358879 to P.K.). The results shown in Fig. 7D, 7E and S7C are based upon data generated by the TCGA Research Network: https://www.cancer.gov/tcga.

## Author contributions

N.P., A.I., J.I. and Z.F. performed the experiments. N.P, A.I., J.H., G.F. and C.D. undertook the data analysis. V.B. provided resources. A.I., J.H., G.F. and C.D. provided methodology. I.B., J.S. and P.K. performed histopathological analysis of CRC specimens. N.P. and J.I. wrote the original draft of the manuscript. J.M., H.H. and A.I. reviewed and edited the manuscript. J.I. and F.J. supervised the project. J.I. and F.J. obtained research funding.

## Material and methods

### Culture and passaging of patient-derived xenografts

#### Animal experiments

Animal experiments performed in France were compliant with French legislation and EU Directive 2010/63. The project was validated by the Ethical Committee (CEEA) no. 26 and was then granted French government authorizations under number 517-2015042114005883 and 2734-2015111711418501. Animal experiments performed in Finland were done in accordance with the Finnish Act on Animal Experimentation (animal license number ESAVI/12558/2021). Mice were obtained from Charles River France and Germany, housed and bred at the Gustave Roussy animal core facility (accreditation number E-94-076-11) and at TCDM (Turku Center for Disease Modelling). Animals were euthanized according to endpoints that were validated by the Ethical Committee, the French government (Ministère de l’Enseignement Supérieur, de la Recherche et de l’Innovation) and the Finnish government.

Three human colorectal tumors (PDX#1 corresponding to LRB-0009C, PDX#2 corresponding to IGR-0012P and PDX#3 corresponding to IGR-014P)) from the CReMEC tumor collection were maintained in NSG mice (strain: NOD.Cg-*Prkdc^scid^ Il2rg^tm1WjI^*/SzJ, female, 6-8 weeks old at grafting) as described in Canet-Jourdan et al. 2022 and in Julien et al. 2012. Briefly, small tumor fragments were subcutaneously engrafted on both flanks of anesthetized mice (2.5% isoflurane). Tumor growth was measured once to twice a week. When the combined tumor burden reached 1700 mm^3^, the mice were euthanized, tumors were used for *ex vivo* experiments and 50 mm^3^ fragments were engrafted on the flanks of new mice in order to maintain the biological material source.

#### Generation of tumoroids

Tumoroids were prepared according to Sato et al., 2011 and adapted for muco-secreting tumors as follows. The PDX#1, PDX#2 or PDX#3 tumors of 1200-1700 mm^3^ were retrieved from the mice, minced into small fragments using a sterile scalpel and were incubated for 1 h at 37°C in a final volume of 5 to 10 ml of culture medium (Dulbecco’s modified Eagle’s medium; DMEM) without fetal bovine serum (FBS) and with 2 mg/ml collagenase (Sigma, C2139). The samples were then mixed with 20 ml of DMEM and filtered on 100 μm mesh size cell strainers (Greiner, EASYstrainer, 542000) and centrifuged 10 min at 277 g. Clusters were isolated from the remaining mucin and single cells by washing in 10 ml of DMEM and pulse centrifugating at 277 g five times. The clusters, now free of mucin and single cells, were maintained for 3 days in ultra-low attachment plates (Corning, CLS3471) in culture medium. Then, organoids were pelleted at 277 g and used for further experiments.

#### Collagen embedding and culture of tumoroids

Collagen-I (Corning, 354236) was neutralized with 1.0 M NaOH and 10× MEM (Life Technologies, 21430-02) according to the ratio: 1.0:0.032:0.1 (v/v/v). The concentration was then adjusted to 2 mg/ml with 1x DMEM, and the collagen-I was incubated on ice for 1h. The organoids, after spending 3 days in suspension as described previously, were then embedded in neutralized collagen-I and were added on top of pre-coated (using 7μl of the collagen mix per well) wells of a μ-Slide 8 Well ibiTreat slide (Ibidi, 80826) at a concentration of 30–50 organoids/5 µl. The gel was allowed to polymerize for 1h at 37°C. Organoids were then cultured in culture medium supplemented with FBS 10% for up to 6 days (3 days for PDX#3). The drugs were diluted in the medium as follows: AIIB2 (DSHB, AB528306, 1 μg/ml), Heregulin-β1 (Peprotech, 100-03-10UG, 20 ng/ml), Trastuzumab (Herceptin, Roche, 10 μg/ml), Pertuzumab (Perjeta, Genentech, 10 μg/ml), Saracatinib (Selleckchem, S1006, 1 μM), EHT-1864 (R&D Systems, 3872, 5 μM), P1E6 (DSHB, AB2619597, 10 μg/ml), FAK14 (Tocris, 3414, 10 μM).

#### Generation of a Patient-Derived Organoid line

The clusters obtained from the PDX as described previously were pulsed centrifuged at 277 g, resuspended in Matrigel (Corning, 354230) and plated in 10 x 15 μl droplets in the bottom of a 6-well plate (Greiner, 657160).

Cells were then incubated at 37°C for 15 minutes to let the basement membrane extract polymerize and culture medium was added as described in Fujii et al 2018, without any human R-spondin1, A83-01 and Afamin-Wnt-3A in serum-free medium. During the first two days, the organoid expansion medium was supplemented with Y-27632 (Calbiochem, 688000, 10 μM). This medium was renewed every two days and PDOs were passaged every 7 to 14 days as described in Cartry et al., 2023.

#### LS513 culture and LS513 TSIPs formation

LS513 cells were obtained from ATCC (#CRL-2134) and cultured in RPMI-1640 medium supplemented with 10% FBS. LS513 were cultured in 10 cm cell culture dish. To generate TSIPs from the LS513 monolayer, the medium was changed every two days until the monolayer reached confluence. After waiting 5 days, the medium was collected and pulse centrifuged at 277g to collect the LS513 TSIPs. These were left for 3 days in suspension in ultra-low attachment plates (Corning, CLS3471), and embedded in collagen as described earlier (using the LS513 medium). For passaging, the monolayer was digested with 1x trypsin when 70% confluence was reached.

#### Immunofluorescence staining

After incubation for 3 days in suspension or for 3 to 6 days in collagen, the apico-basolateral polarity of organoids was quantified. Cells were washed thrice with PBSCM (PBS supplemented with CaCl_2_ (0.1 mM) and MgCl_2_ (1 mM)), fixed with 4% paraformaldehyde (PFA) for 5 minutes (for spheres in suspension) or 45 minutes (for spheres in collagen) at RT. Spheres fixed in suspension were then embedded in collagen for imaging as previously described. Permeabilization was then performed in PBSCM supplemented with 0.5% Triton X-100 for 45 minutes. Spheres were incubated with primary antibodies overnight at 4°C with the dilutions mentioned in Table 2 in PBSCM supplemented with 10% FBS and 0.1% Triton X-100. After washing thrice with PBSCM supplemented with 0.05% Tween, spheres were incubated with secondary antibodies and phalloidin for 2h at RT with the dilutions mentioned in Table 2, as well as with DAPI (1 μg/ml). Spheres were then washed thrice with PBSCM supplemented with 0.05% Tween. The gel was then immerged in PBS before imaging.

**Table 2.**
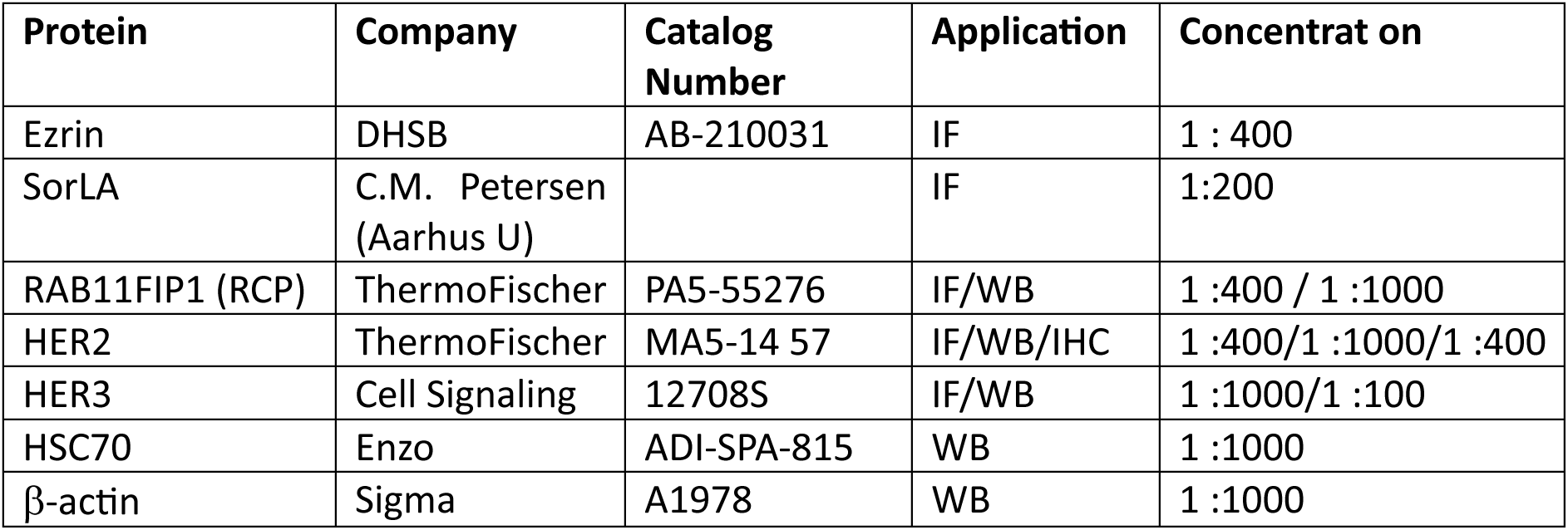
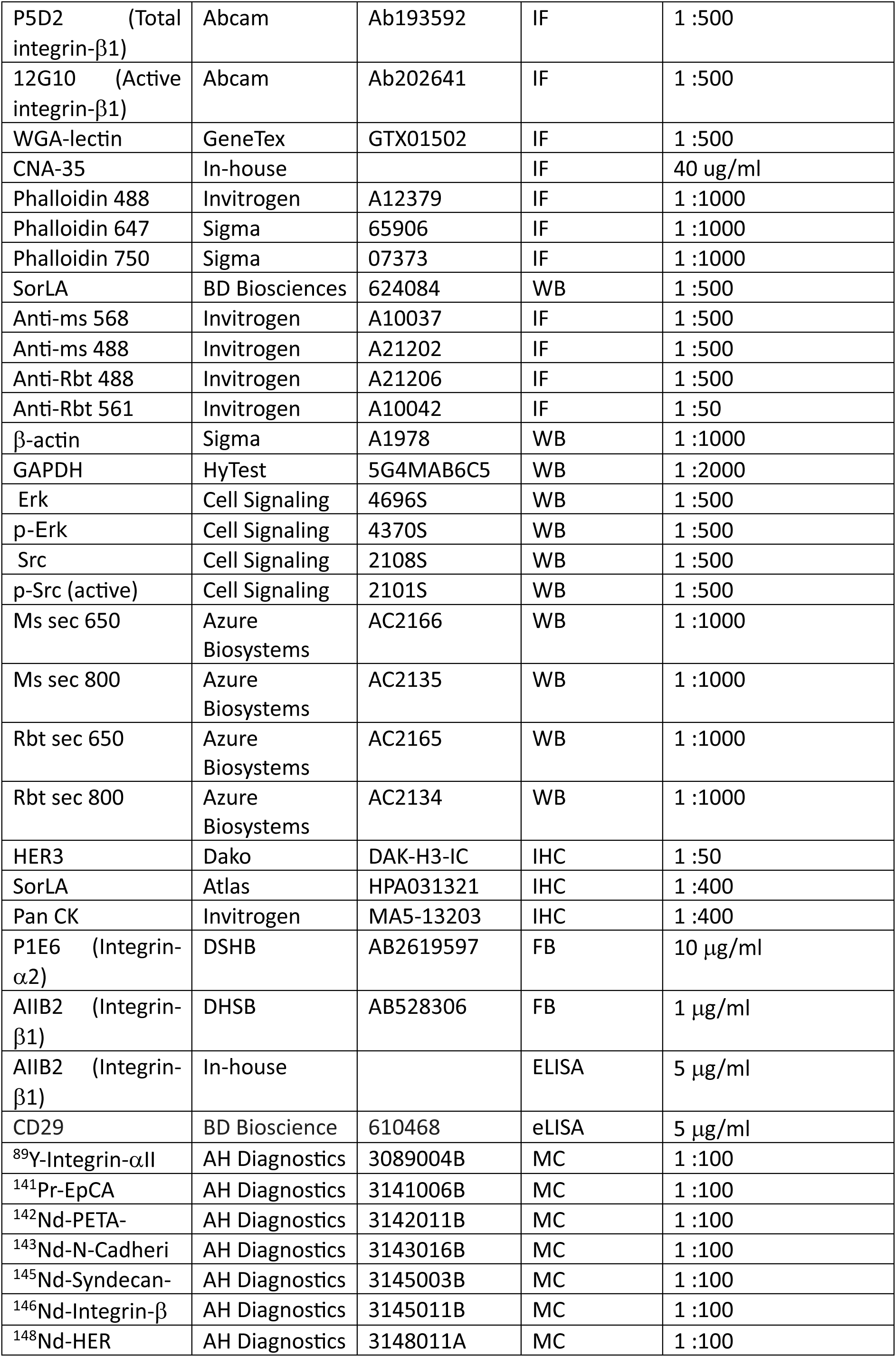
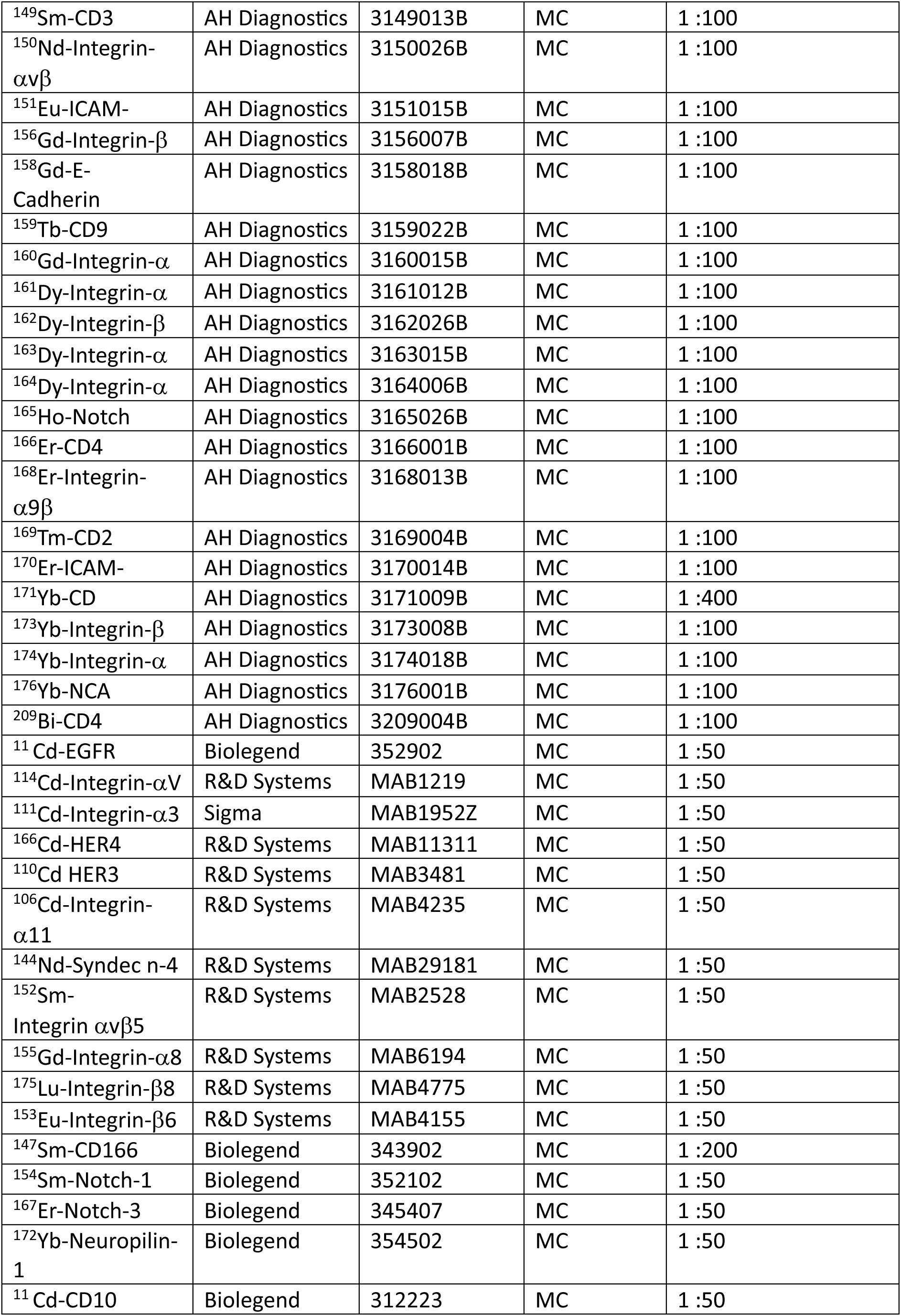
– Antibodies and Reagents (WB=Western Blot, IF=Immunofluorescence, IHC=Immunohistochemistry, FB=Function blocking, MC=Mass cytometry, ELISA=enzyme-linked immunosorbent assay)

#### Confocal imaging

Images were acquired either using a SpinningDisk CSU-W1 microscope (Yokogawa) with a ZylasCMOC camera piloted with an Olympus X83, or with a 3i CSU-W1 spinning disk confocal microscope with Hamamatsu ORCA-Flash4.0 v2 scientific complementary metal-oxide semiconductor (sCMOS) camera (x40 water immersion objective, 1.1 NA). For traction force microscopy experiments and live imaging, a x20 objective was used. Basic image processing including look up table optimization, channels overlay, projections, etc… were performed using Fiji software (Schindelin et al., 2012). More specific quantifications are detailed in the following sections. In panels showing Ezrin staining to assess the polarity phenotype, the signal balance was adjusted for a better visualization. These adjustments were only done on representative shown images and not on the image data used for quantitative analysis.

#### Western-blotting

Spheres embedded in collagen were first released from the matrix by incubation in DMEM without serum supplemented with 2 mg/ml collagenase (Sigma, C2139) for 45 minutes. After pulse centrifugating at 277g, spheres were washed with PBS and pulse centrifugated at 277 g twice. Spheres were then lysed in TXLB buffer [50 mM Hepes, 1% Triton X-100, 0.5% sodium deoxycholate, 0.1% SDS, 0.5 mM EDTA, 50 mM NaF, 10 mM Na_3_VO_4_, and protease inhibitor cocktail (cOmplete Mini, EDTA-free, Roche)]. Cells cultured in 2D were washed twice with PBS and directly lysed with TXLB. Separation was performed by gel electrophoresis (Mini-PROTEAN TGX Precast Gels 4-20%, Bio-Rad, 4561096), before transferring onto a nitrocellulose membrane (Trans-Blot Turbo Transfer System, Bio-Rad) and blocking with AdvanBlock-Fluor (Advansta, R-03729-E10). Primary antibodies in AdvanBlock-Fluor were incubated overnight at 4°C with the dilutions mentioned in Table 2. Membranes were washed thrice between primary and secondary antibody treatments with Tris-buffered saline with 0.1% Tween 20 (TBST). IRDye secondary antibodies (see Table 2) were incubated for at least 1 hour at RT, before detection on an Odyssey fluorescence imager CLx (LI-COR). Densitometry analysis was performed in Fiji by normalizing the signal to a suitable loading control (either GAPDH, β-actin or HSC70).

#### PNGase digestion of lysates

For PNGase digestion, cell lysates were prepared in SDS-free buffer [50 mM Hepes, 1%NP-40, 0.5% sodium deoxycholate, 0.5 mM EDTA, 50 mM NaF, 10 mM Na_3_VO_4_, and protease inhibitor cocktail (cOmplete Mini, EDTA-free, Roche)]. 9 μl of lysate was mixed to 1 μl of Glycoprotein Denaturing Buffer 10X) (NEB, B1704S). The lysate was denatured at 100°C for 10 minutes, then chilled on ice. 2 μl GlycoBuffer 2 (10X) (NEB, B3704S), 2 μl 10% NP-40 (NEB, B2704S), 5 μl H_2_O and 1 μl PNGase F (NEB, P0704S) were then added to the lysate, and mixed gently. The lysate was then incubated at 37°C for 1h. From there, the samples were prepared and run as described before.

#### SorLA silencing using shRNA lentiviral transduction

During passaging of the PDX#3-generated PDO line as described previously, 10^5^ cells were resuspended in 36 μl of organoid expansion medium and infected with 4M TU [62 μl of each virus (Origene, TL309181V)], 1 μl polybrene (Merck, TR-1003-G, stock solution at 1 mg/ml) and 1 μl Y-27362 (Calbiochem, 688000, 10 μM, stock solution 1 mM), in a U-bottom 96-well plate (Falcon, 351177) for 6 hours at 37°C. The content of the wells were then collected in a 1.5 mL Eppendorf tube, 1 ml of DMEM+10%FBS was added, and cells were centrifugated at 200 g for 3 minutes. Cells were then resuspended in 60 μl Matrigel (Corning, 354230) and 4×15 μl droplets were poured in a 24-well cell culture plate (Cellstar, 662 160) and left at 37°C degrees to polymerize for 15 minutes. 1 ml organoid expansion medium was used for organoid expansion (supplemented with 10 μM Y-27362 for the first two days). The medium was changed every two days and organoids were left to grow for 7 days. For polarity assays, the well was first washed thrice with PBS, before adding 1 ml of Cell Recovery Solution (Corning 354253). Cells were then left to incubate at 4°C for 20 minutes, and mechanical dissociation was performed with a p1000 pipette until Matrigel was completely dissolved. 4 ml of PBS was added and spheroids were pulse centrifuged at 277 g. The spheres, now free of Matrigel, were maintained for 3 days in ultra-low attachment plates (Corning, CLS3471) in PDX culture medium. Then, organoids were pelleted at 277 g and used for further experiments.

#### SORLA silencing using siRNA transient transfection

LS513 were plated in a 6 well plate at 80% confluency. Transient siRNA transfections were performed using Lipofectamine RNAiMAX reagent (Invitrogen, 56532) according to the manufacturer’s instructions. SORLA-targeting siRNAs were ON-TARGETplus obtained from Dharmacon—siSORLA #1 (J-004722-08), siSORLA #2 (J-004722-06), siSORLA #3 (J-004722-07), siSORLA #4 (J-004722-05). For controls, Allstars negative control (Qiagen, Cat. No. 1027281) was used. siRNA concentrations used were all 20 nM and cells were transfected with siRNAs 72 h prior to experiments.

#### Integrin uptake assay

Surface biotinylation-based integrin trafficking assays in SorLA-silenced LS513 cells were performed based on previously published methods (Farage et al., 2021; Arjonen et al. 2012), with some modifications. Nunc MaxiSorb 96-well plates (Thermo Fischer, 44-2404-21) Enzyme-linked immunosorbent assay (ELISA) plates were coated with anti-β1-integrin antibody mix (5 μg/ml of AIIB2 (in-house) and anti-CD29 (BD Bioscience #610468)) in TBS (50 μl per well) overnight at +4 °C. The wells were blocked with 5% BSA in TBS for 2 h at 37 °C (100 μl per well). LS513 cells were silenced three days before the experiment as described earlier. 2 hours prior to the experiment, the medium was changed to prewarmed RPMI with 10% FBS to induce receptor traffic. The cells were placed on ice and washed once with cold PBS. Cell surface proteins were labelled with 0.13 mg/ml EZ-link cleavable sulfo-NHS-SS-biotin (Thermo Scientific, 21331) in serum-free RPMI medium for 30 min at +4 °C. Any unbound biotin was removed by washing with cold medium and pre-warmed RPMI+10% FBS with or without 100 μM primaquine (Sigma, 160393) was added to the cells. The biotin-labelled surface proteins were allowed to traffic at +37 °C for 15 or 30 minutes. Cells were placed on ice, washed once with cold PBS and cold cell surface reduction buffer (50 mm Tris– HCl, pH 8.6 and 100 mm NaCl). Cell surface biotin was cleaved with non-membrane permeant reducing reagent MesNa (30 mg/ml, sodium 2-mercaptoethanesulfonate; Fluka, 63705) in cell surface reduction buffer 20 min at 4 °C, followed by quenching with 100 mM iodoacetamide (Sigma, I3750) for 15 min on ice. For the 0 min internalization, cells were maintained on ice in serum-free RPMI until cell surface reduction with MesNA. The cells were lysed by scraping in lysis buffer (1.5% octylglucoside, 1% NP-40, 0.5% BSA, 1 mM EDTA, and protease and phosphatase inhibitors) and incubation at +4 °C for 20 min with rotation and cleared by centrifugation (16,000*g*, 10 min, 4 °C). To detect the biotinylated integrins, 50 μl volumes of the cell lysates were incubated in duplicate wells at +4 °C overnight, washed extensively with TBST, incubated for 2 h at 4 °C with 1:1,000 horseradish peroxidase-coupled streptavidin (Fisher, 21130), washed and detected with antibody for ELISA detection.

#### Immunohistochemistry

Histological CRC and peritoneum specimens obtained after surgical resection were formalin fixed and paraffin embedded according to routine protocols. Sections (3 μm) of formalin-fixed and paraffin-embedded samples were deparaffinized, unmasked (pH 8) and rehydrated before immunohistochemistry or immunofluorescence stainings.

Immunohistochemistry sections were immunostained for SorLA, HER2 and HER3 (see Table 2). Slides were imaged using Axioscan Z1, Zeiss (x20, NA 0.8). Due to differences in the three different stains, imaging was optimized separately for each channel, but the settings were then kept constant for all samples. Regions of interest (ROIs) were hand-drawn using QuPath (Bankhead et al., 2017), exported as TIFF files through custom scripting, and then stored as Zarr datasets. These were subsequently fed into a custom processing pipeline using the Python scientific library scikit-image (Van der Walt et al., 2014). Zone binary masks for apical-in and apical-out ROIs were generated using digital image processing. For apical-out ROIs, zero-intensity pixels were used to delineate the mask boundary; for apical-in ROIs, a combination of Gaussian filter, local thresholding and binary morphological operations was used, with masks drawn manually when required. Within the masked zones, nuclear counts based on the DAPI channel were generated using a StarDist (Schmidt et al., 2018; Weigert et al., 2022) pre-trained model “2d_versatile_fluo”. The total intensity signal for SorLA, HER2, and HER3 were measured in apical-in and apical-out ROIs and normalized against the nuclear count in respective ROIs. Pearson’s test was used for correlation analysis between HER2, HER3 and SorLA. The complete source code repository for image processing is included in the supplemental materials.

#### Mass cytometry

Spheroids grown in collagen for 3/6 days underwent collagen digestion by incubation in DMEM without serum supplemented with 2 mg/ml collagenase (Sigma, C2139) for 45 minutes. After resuspending in DMEM + 10% FBS and pulse centrifugating at 277 g, they were collected. Spheroids cultured in suspension, clusters obtained from PDX digestion or spheroids released from collagen as described previously were washed with PBS thrice and pulsed centrifuged at 277 g thrice in order to keep the clusters/spheroids and get rid of any single cells and secreted mucins. The spheres were then digested in Cell Dissociation Buffer Enzyme-Free PBS-based (Gibco, 13151-014) supplemented with 2 mg/ml collagenase (Sigma, C2139) and incubated for 1h at 37°C with occasional mechanical dissociation by pipetting. After addition of DMEM+10% FBS to quench the collagenase, cells were centrifuged at 200 g for 3 minutes and washed thrice with PBS. They were then resuspended in 1 ml cold PBS and filtered through a 5 ml polystyrene round bottom tube with Cell-Strainer cap (Falcon, 352235), and kept on ice until staining, which was done according to the Fluidigm Maxpar Cell Surface Staining with Fresh Fix protocol. The sample was then run using a Helios™ Mass Cytometer and the data were analyzed with Cytobank and clustered/visualized through SPADE and tSNE analysis.

#### Generation of fluorescent collagen

To fluorescently label rat tail type I collagen (∼4.24 mg/mL, 354236, Corning), 1.65 mL was mixed with 450 μL of Milli-Q water and 500 μL of neutralizing buffer (20 mM NaH2PO4, 112 mM Na2HPO4⋅2H2O, 0.4 M NaCl, and 46 mM NaOH) and incubated at 37 °C for 30 min. The polymerized collagen was then washed thrice with PBS for 10 min. Then, 3 mL of Milli-Q water and 1 mL of bicarbonate buffer [1 M NaHCO3 (pH 8), raised dropwise to pH 8.3 using 1.17 M Na2CO3 (pH 11)] were added to the collagen gel before addition of the Alexa Fluor™ 647 NHS Ester (Succinimidyl Ester) dye (A20006, Invitrogen) in 100 μL of PBS. After incubating the collagen mix overnight at 4 °C, the dye was then removed, and the collagen was washed with PBS with rotation at room temperature for 30 min, changing the PBS five times. Stained collagen was then depolymerized through the addition of 2 mL HCl (20 mM) and gentle rotation at 4 °C overnight. The collagen was finally centrifuged at 20,000 g for 10 min, collecting the labeled collagen from the supernatant. The fluorescent collagen was then used at a 1:1000 concentration in the previously described neutralized collagen gel.

#### Cloning and generation of CNA35

To generate the CNA35-mScarlet construct, pET28a-CNA35-EGFP (A kind gift from Maarten Merkx (Eindhoven University of Technology, MB Eindhoven, The Netherlands), Addgene plasmid #61603) was digested with NheI/EcoRI and ligated with a NheI/EcoRI digested mScarlet gene fragment (IDT) to generate pET28a-mScarlet-CNA35. The final construct was validated by analytical digestion and sequencing. Recombinant protein production and purification for CNA35-mScarlet was performed as described previously (Conway et al., 2023).

#### *Ex vivo* peritoneum assay

Peritoneum samples were collected from mice and decellularized by incubating them in a 1M NH_4_OH solution for 1h at RT. After washing thrice with PBS for 15 minutes, peritoneum samples were left to incubate with PBS and penicillin-streptomycin (1:100) at 4°C overnight. After washing thrice with PBS for 15 minutes, the peritoneum was sectioned into 1cm x 1cm pieces and adhered (using Tissue Adhesive, 3M, 1469SB) to the bottoms of plastic Transwell inserts (Greiner, Thincerts, 8 um pore size, 662638) after removing the filter with a scalpel. 100 tumor spheres were resuspended in 100 ml of DMEM+10%FBS and placed in the well with AIIB2 (DSHB, AB528306, 1 μg/ml), Y27632 (Calbiochem, 688000, 25 μM), Heregulin-β1 (Peprotech, 100-03-10UG, 20 ng/ml), Trastuzumab (Herceptin, Roche, 10 μg/ml), and/or Pertuzumab (Perjeta, Genentech, 10 μg/ml) for 6 days. The fixing and IF staining was performed as described previously. Peritoneum bits were placed upside-down on a glass bottom dishes (Cellvis, D35-14-1-N), and imaged as described previously, using the x20 objective.

#### Whole Exome Sequencing

DNA was extracted using the DNeasy kit (Qiagen, Cat. No. 69504) from organoids either after 3 days in suspension (washed once in PBS supplemented with Ca^2+^ and Mg^2+^ as mentioned above). Whole exome analysis was performed by Integragen SA (France), comparing the three samples (PDX#1, PDX#2 and PDX#3) to a PON (panel of normal), and analyzed with MERCURY™.

#### Collagen orientation analysis

Type I collagen gels with PDX#3 and PDX#2 spheroids were prepared on glass bottom dishes (Cellvis, D35-14-1-N). 80 µl of PureCol EZ Gel (Advanced BioMatrix, 5074) was spread on the glass bottom using a micropipette tip and allowed to polymerize at +37 °C for 1 h. Next, PDX#3 or PDX#2 spheroids were pulse centrifuged at 277 g to remove the mucin and single cells. Approximately 100 spheroids were mixed with 80 µl of PureCol EZ Gel and pipetted on top of the previously polymerized collagen layer, after which the mixture was allowed to polymerize at +37 °C for 1 h. Spheres were treated with AIIB2 (DSHB, AB528306, 1 μg/ml), Y27632 (Calbiochem, 688000, 25 μM), Heregulin-β1 (Peprotech, 100-03-10UG, 20 ng/ml), Trastuzumab (Herceptin, Roche, 10 μg/ml) and/or Pertuzumab (Perjeta, Genentech, 10 μg/ml). One day before the samples were imaged, the cultures were supplemented with 1:1000 SiR-actin (Sprichrome, SC001) and ∼40 µg/ml of mScarlet-conjugated collagen probe CNA35 (Aper et al., 2014) (see above).

The spheroids were imaged live using a Marianas spinning disk confocal microscope, 20x objective, Orca Flash4 sCMOS camera, and 2×2 binning (see Microscopy for details) at 37°C, 5% CO_2_. 60-80 µm stacks were acquired around the center (z) of each spheroid. In order to analyze collagen fiber orientation around the spheroids, ∼4 µm substacks were acquired near the center of each spheroid and used for creating maximum intensity projections. Next, 200 x 200 µm regions of interest depicting CNA35 directly proximal to each spheroid (but excluding any dense collagen aggregates on the spheroid surface) were selected from the projections for analysis. If the matrix surrounding the spheroid was obviously heterogeneous, the region was selected to maximize the local alignment.

The selected regions were analyzed with ImageJ plugin OrientationJ, using cubic spline gradient and a local window size of 4 pixels. In the color survey, hue represented orientation and saturation represented coherency. All the local orientations were exported and analyzed in R script to yield fiber orientation indices (Ferdman et al., 1993; Taufalele et al., 2019). Briefly, the orientations (−90°…+90°) representing each region were normalized, i.e., their distribution was centered around zero based on the peak of the histogram. Next, orientation indices (*S*) were calculated such that

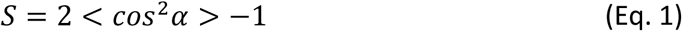

where *α* is the angle between an individual (fiber) orientation and the average orientation of the entire region, and <*cos^2^α*> is the average square cosine of all *α* per analyzed region. And index of 0 represents a random distribution, and an index of 1 represents a perfectly aligned distribution.

#### Collagen displacement fields

In order to measure transient displacements exerted on the collagen matrix by the PDX#3 and PDX#2 spheroids, the spheroids were prepared and embedded in type I collagen, as described above. The spheroids were grown in the gels for 6 days and supplemented with 1:1000 SiR-actin and ∼40 µg/ml mScarlet-CNA35 one day before the imaging. 60-80 µm stacks were acquired around/near the center of each spheroid, before and after the cells and matrix were relaxed by adding 10 µM latrunculin B and incubating for 20 minutes. Marianas spinning disk confocal microscope, 20x objective, Orca Flash4 sCMOS camera, and 2×2 binning were used for the imaging (see Microscopy for details).

3D displacement fields were calculated using TFMLAB (Barrasa-Fano et al., 2021a; Sanz-Herrera et al., 2023; Barrasa-Fano et al., 2021b), a traction force microscopy toolbox implemented in MATLAB R2022a (MathWorks). The spheroids were segmented using actin images, variable threshold adjustment and a minimum object size of 10^4^ voxels. Rigid image registration was conducted using the default phase correlation-based algorithm. The displacements were calculated from CNA35 images using 10×10×10 µm grid spacing, default registration metric and optimizer (normalized correlation coefficient, adaptive stochastic gradient descent) and post-shift correction. The results were visualized using ParaView v5.11.0 (Ahrens et al., 2005).

#### Analysis of SORLA and ITGB1 gene expression in human tumors

Preprocessed TCGA colon adenocarcinoma cohort RNAseq data and raw RSEM-counts were downloaded from https://gdc.cancer.gov/node/905/ and https://gdac.broadinstitute.org/runs/stddata 2016_01_28/data/, respectively. CMSCaller (Eide et al., 2017) was used to infer consensus molecular subtypes (CMS) from RSEM-counts, excluding calls with FDR > 0.05. The samples were categorized as mucinous and control cases based on previously conducted characterization (Nguyen et al., 2021). Associations between SORLA and ITGB1 gene expression were assessed with the preprocessed normalized log2 mRNA expression data by computing linear regressions within each CMS group.

#### Statistical analysis

All statistical comparisons were performed using Prism 7 (GraphPad software), as indicated in the figure legends, repeating all experiments independently at least three times.

