## Supplementary Material for "Signaling downstream of tumor-stroma interaction regulates mucinous colorectal adenocarcinoma apicobasal polarity"

Supplementary information

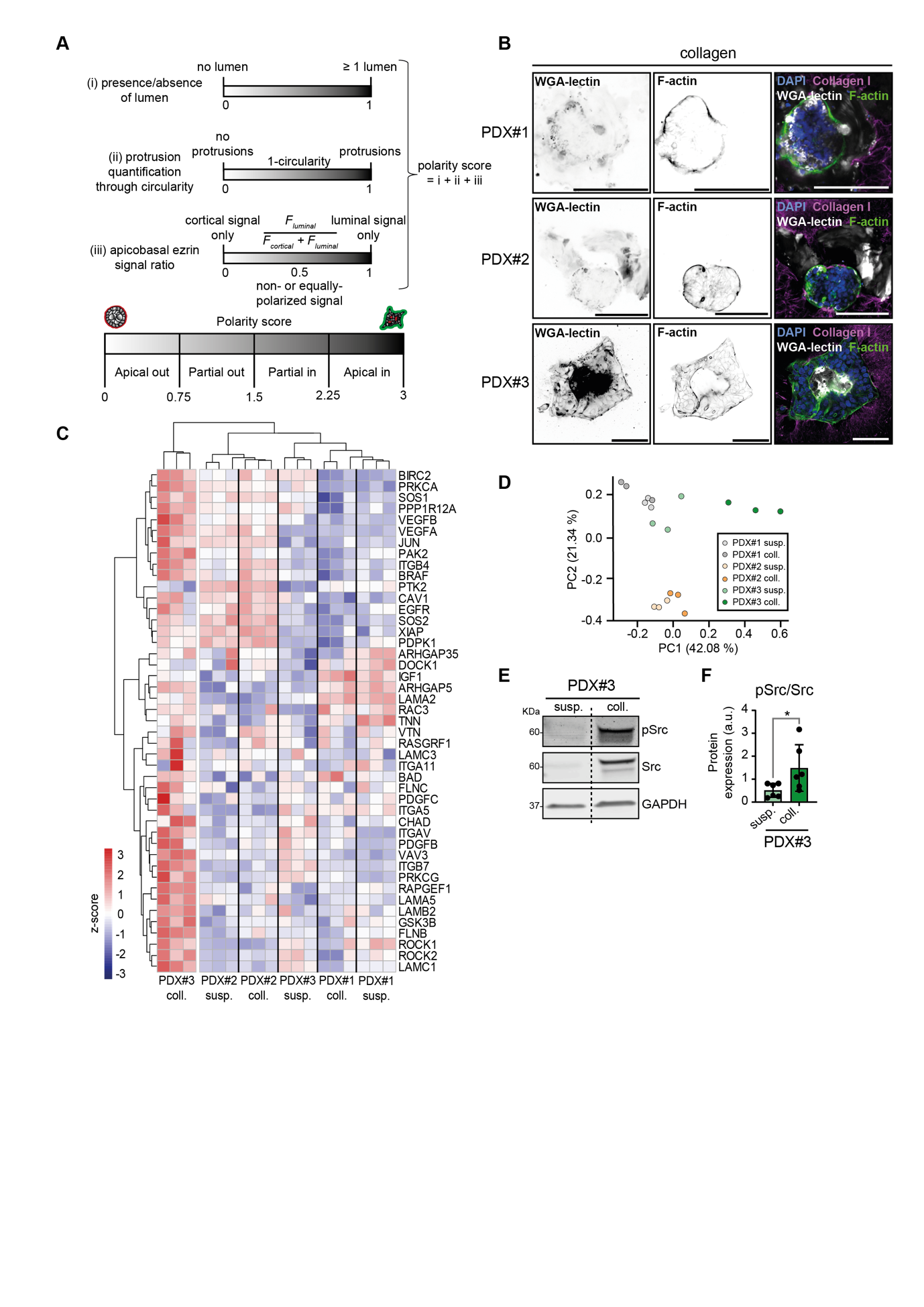
**FIGS1. Characterization of apicobasal polarity of mucinous CRC PDX models**

(A) Outline of polarity score calculation used for all experiments. The score (from 0: apical-out to 3: apical-in) is a sum of three parameters, i) the presence of a lumen, ii) circularity, and iii) ratio of luminal/cortical Ezrin fluorescence. (B) Representative immunofluorescence images of PDX#1-, PDX#2- and PDX#3-generated tumoroids in collagen, with labeled F-actin and WGA-lectin to detect mucin secretion. (C) Heatmap of the 45 top differentially regulated genes in the KEGG Focal Adhesion Pathway in PDX#3 (collagen vs. suspension). (D) Associated Principal Component Analysis on the same geneset for PDX#1, PDX#2 and PDX#3 models in suspension and collagen. (E, F) Western blot analysis of pSrc and Src levels in PDX#3 tumoroids in suspension or collagen and associated quantifications (n = 6, paired t-test, p=0.0361). Scale bars, 100 μm.

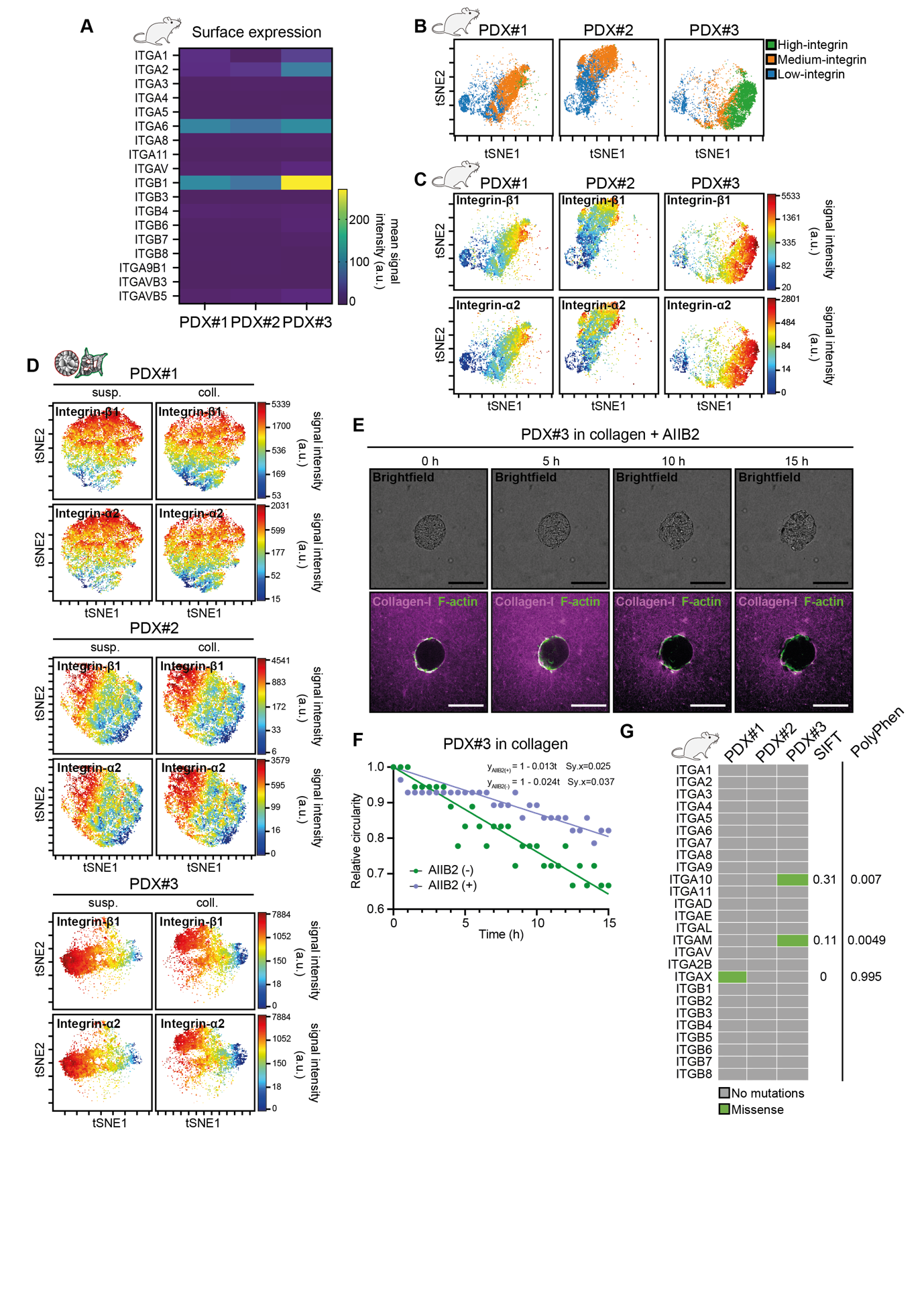

**FIGS2. Integrin surface expressions in PDX#1, PDX#2 and PDX#3 tumors and generated tumoroids**

(A) Heatmaps representing mean cell-surface expression of the indicated integrins in PDX#1, PDX#2 and PDX#3 tumors dissected from mice. (B) FlowSOM clustering of integrins following a tSNE analysis of single-cell mass cytometry data obtained from PDX#1, PDX#2 and PDX#3 tumors out of mice shows three distinct cell clusters based on integrin surface expression levels. (C) tSNE analysis of single-cell mass cytometry data obtained from dissected PDX#1, PDX#2 and PDX#3 tumors based on the whole panel of surface receptors (see Material and Methods, table 2), overlayed with the intensity of integrin-α2 and integrin-β1 signal intensities. (D) tSNE analysis of PDX#1-, PDX#2- and PDX#3-generated tumoroids in suspension and collagen based on the whole panel of surface receptors, overlayed with the intensity of integrin-α2 and integrin-β1 signal intensities. (E) Snapshots of a live polarity reversion of PDX#3 over 15 hours, with AIIB2 (1 μg/ml), starting from the embedding in collagen timepoint (Control in Fig. 1C). (F) Quantification of the sphere circularity of **Videos 1** and **2**, showing different evolutions when the spheres are cultured with or without AIIB2. (G) Whole-Exome Sequencing data, showing mutation profiles in PDX#1, PDX#2 and PDX#3 for the different integrin subunits. Scale bars, 100 μm.

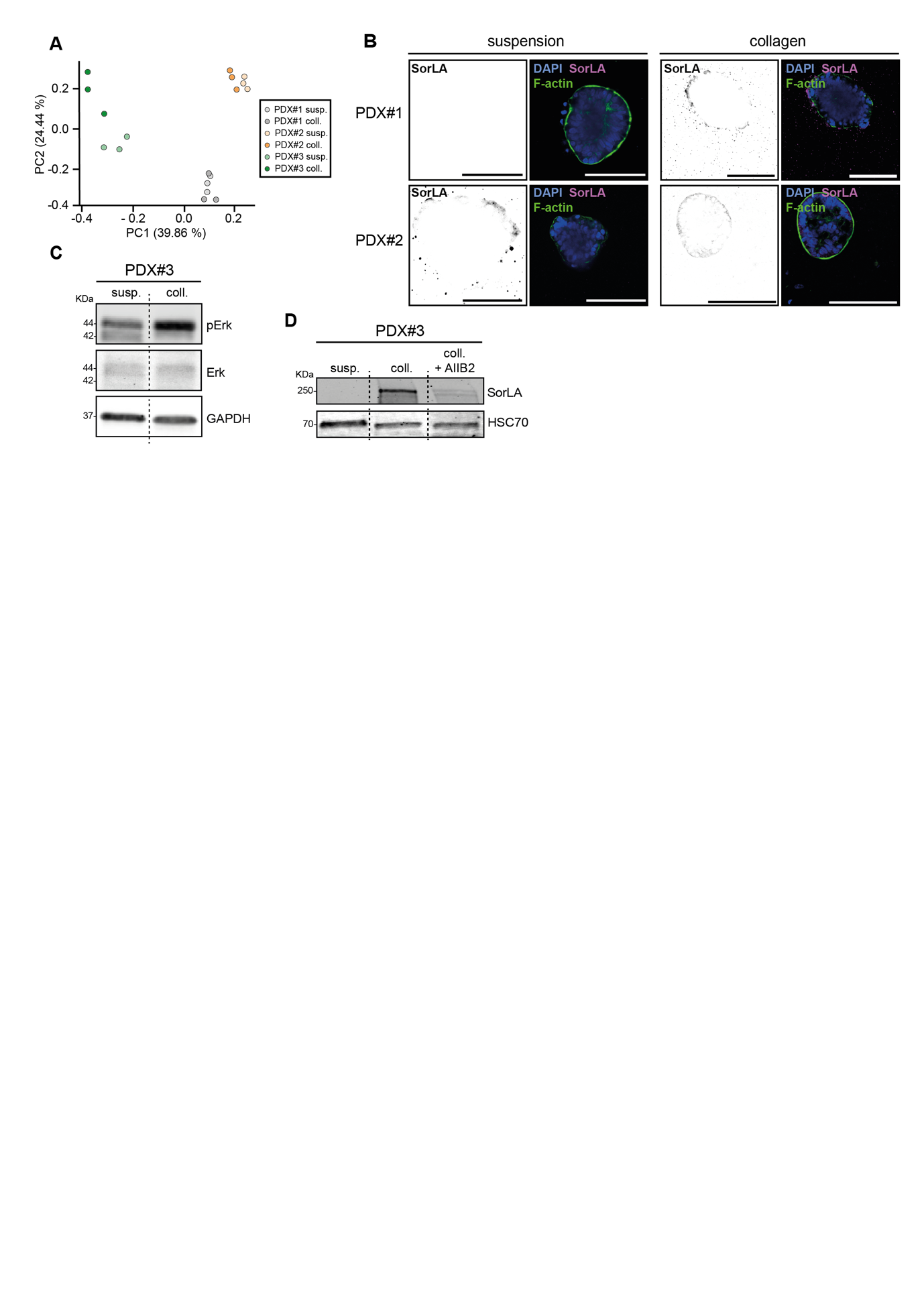

**FIGS3. SorLA signaling regulates polarity in MUC CRCs**

(A) PCA associated to the relative expression of key regulators in integrin-β1 trafficking, generated from an RNA microarray analysis of PDX#1-, PDX#2- and PDX#3-generated tumoroids in suspension and in collagen. (B) Immunofluorescence of PDX#1- and PDX#2-generated tumoroids in suspension and in collagen, showing SorLA staining. (C) Western-Blot of pErk, Erk and GAPDH as a loading control in PDX#3-generated tumoroids in suspension and collagen. (D) Western-blot of SorLA and HSC70 as loading control in PDX#3-generated tumoroids in suspension, collagen, and collagen + AIIB2 (1 μg/ml). Scale bars, 100 μm.

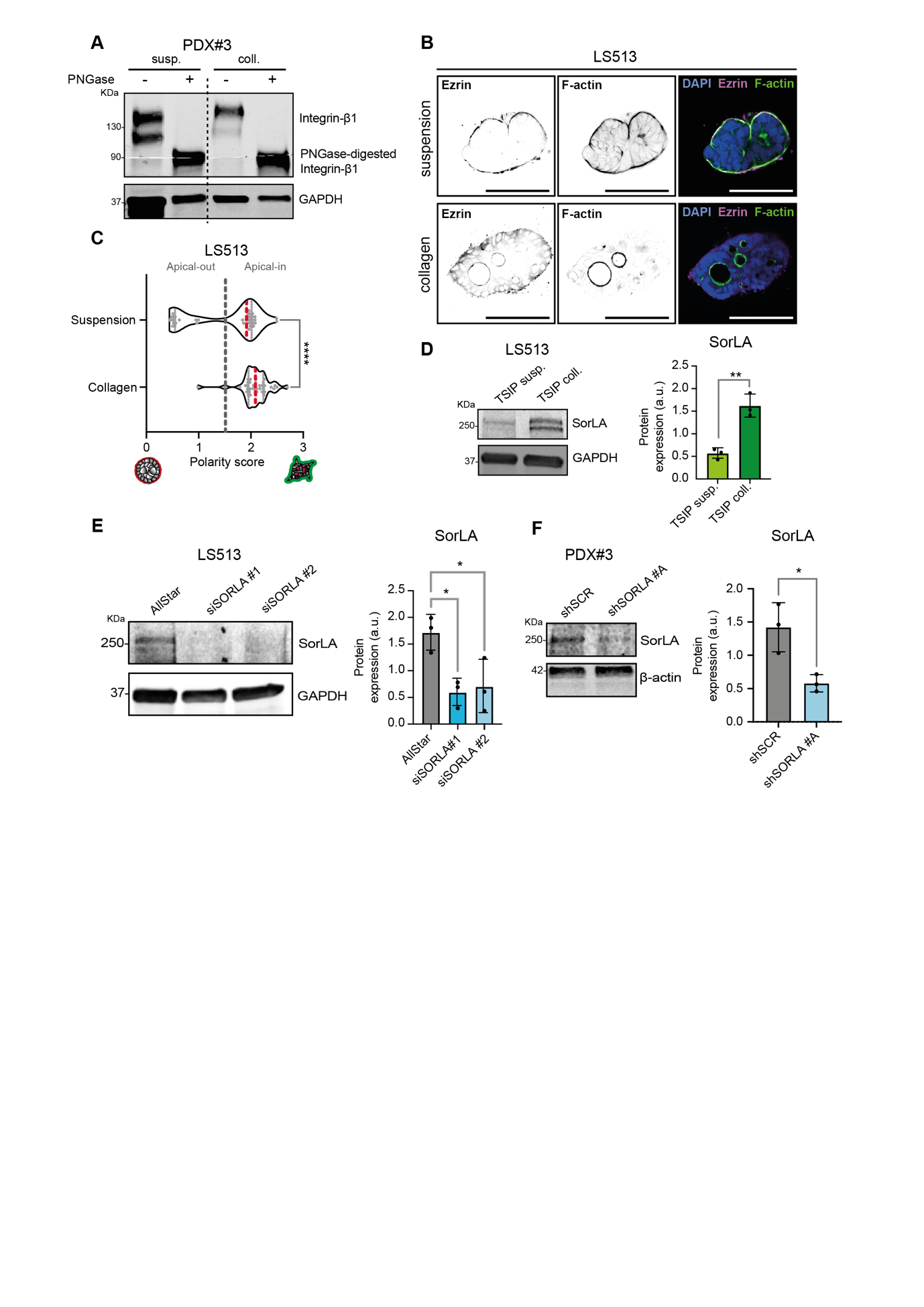
**FIGS4. SorLA-dependent Integrin-β1 recycling loop in PDX#3.**

(A) Western-Blot of integrin-β1 in PDX#3-generated tumoroids in suspension and collagen, before and after PNGase digestion of the lysates. (B) Immunofluorescence of LS513 TSIPs in suspension and in collagen, showing their polarity status through Ezrin staining. (C) Comparison of polarity scores amongst models in collagen (n_suspension_=42, n_collagen_=40, Mann-Whitney test, p-value<0.0001). (D) Western-blot of SorLA in LS513 TSIPs in suspension and collagen (n=3, t-test, p-value=0.0029). (E) Western-blot of SorLA in LS513 transfected with SORLA-targeting siRNA (n=3, ANOVA, p-value_#1_=0.0259, p-value_#1_=0.0393). (F) Western-blot of SorLA in PDX#3-generated organoids infected with SORLA-targeting shRNA (n=3, t-test, p-value=0.0203). Scale bars, 100 μm.

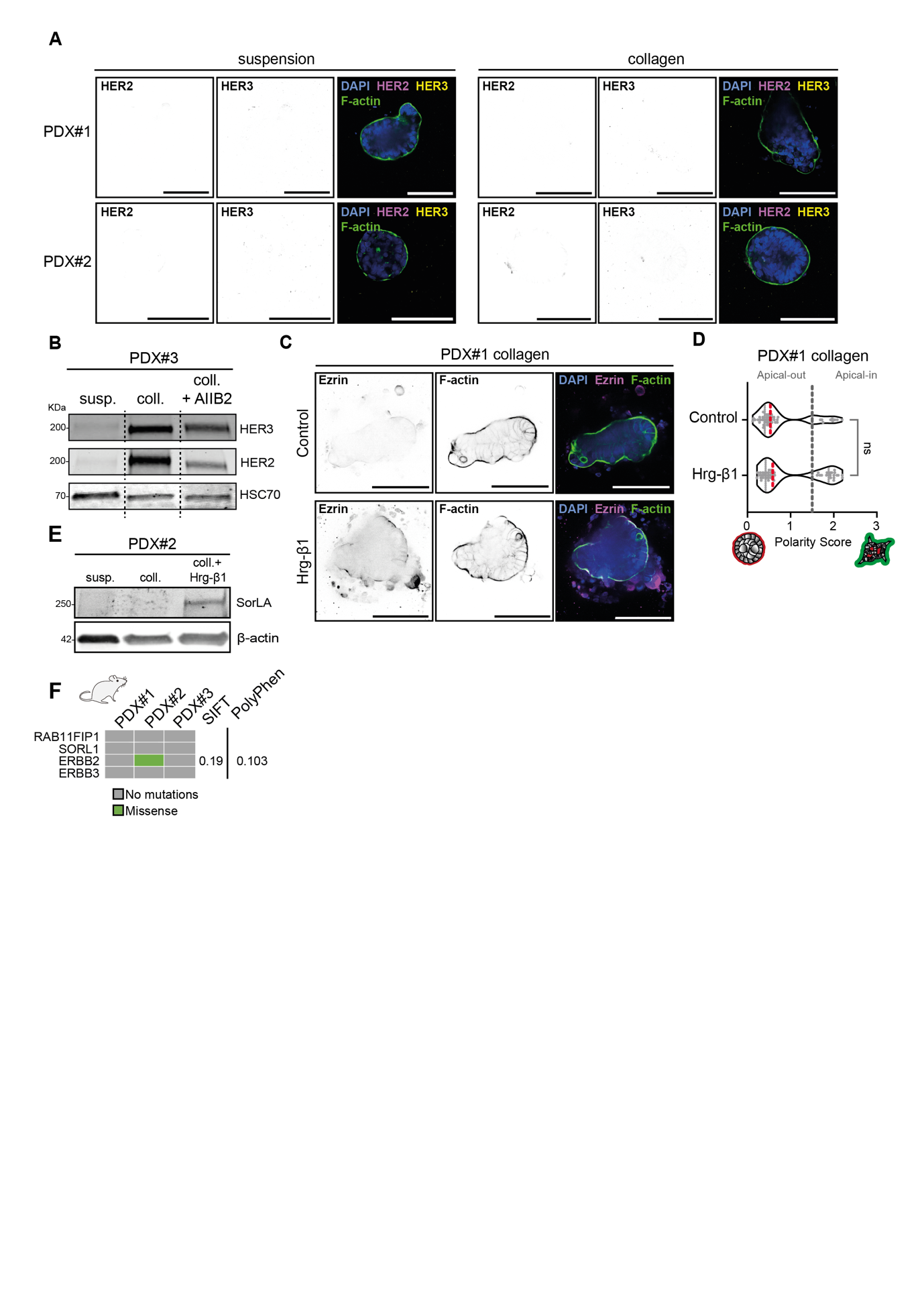

**FIGS5. HER2 and HER3 characterization in MUC CRC PDXs.**

(A) Immunofluorescence of HER2 and HER3 in PDX#1- and PDX#2-generated tumoroids in suspension and in collagen. (B) Western-Blot showing HER2 and HER3 expression in PDX#3-generated tumoroids in suspension, collagen, and collagen + AIIB2 (1 μg/ml). (C) Immunofluorescence of PDX#1-generated tumoroids in collagen, showing their polarity status through Ezrin staining after Hrg-β1 treatment (20 ng/ml). (D) Comparison of polarity scores in PDX#1 in collagen after treatment with Hrg-β1 treatment (20 ng/ml). (n_Ctrl_=59, n_Hrg-β1_ =55), Mann-Whitney test, p=0.2359). (E) Western-blot showing SorLA expression in PDX#2-generated tumoroids in suspension, collagen and collagen + Hrg-β1. (F) Whole-Exome Sequencing data, showing mutation profiles in PDX#1, PDX#2 and PDX#3 in *RAB11FIP1*, *SORL1*, *ERBB2* and *ERBB3*. Scale bars, 100 μm.

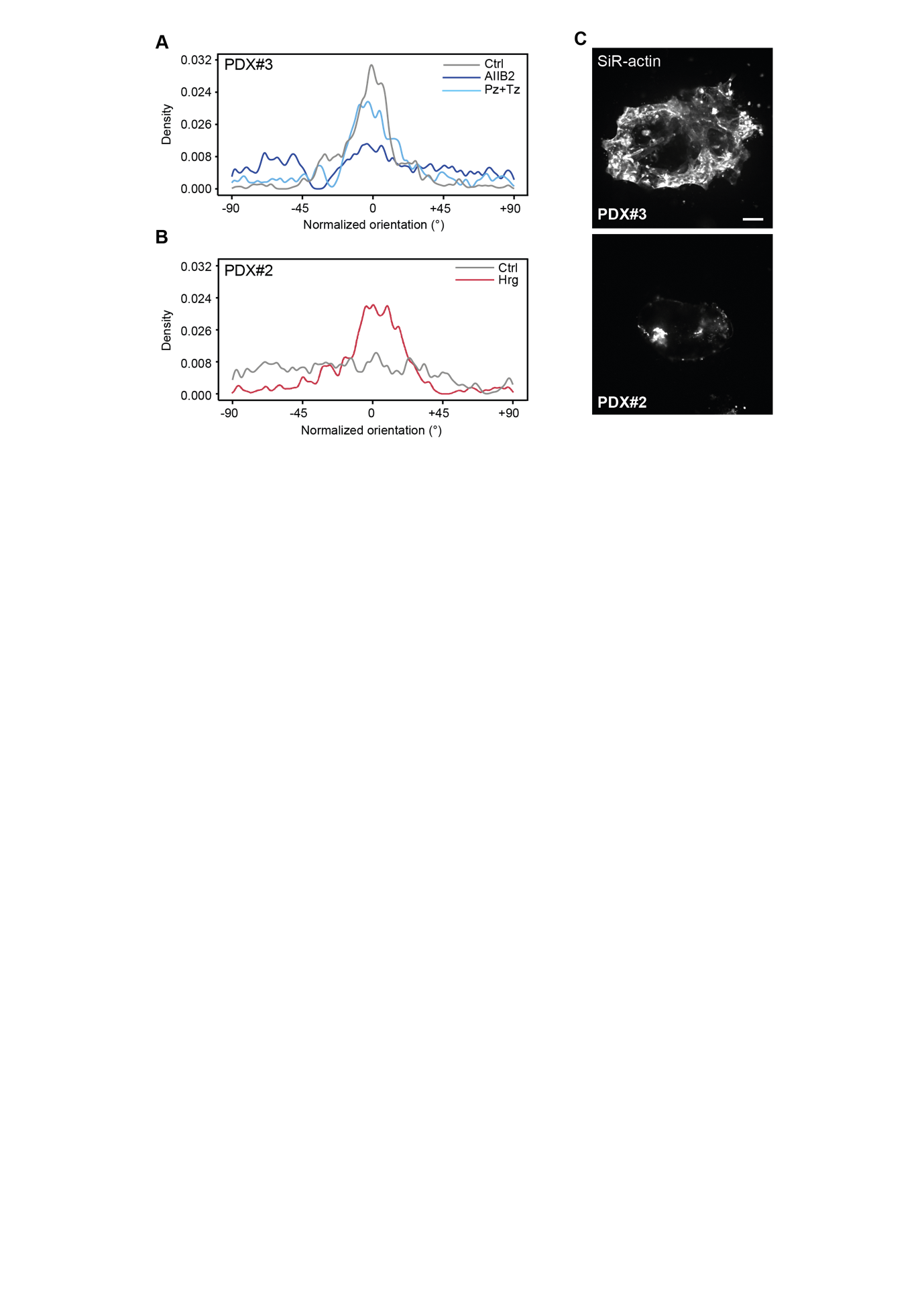

**FIGS6. Local collagen alignment proximal to PDX#2 and PDX#3 tumoroids.**

(A) Normalized orientation of collagen fibers proximal to PDX#3-generated tumoroids after 6 days in collagen, with AIIB2 treatment (1 μg/ml) or Trastuzumab+Pertuzumab treatment (Pz+Tz, 10 μg/ml, 10 μg/ml) (B) Normalized orientation of collagen fibers proximal to PDX#2-generated tumoroids after 6 days in collagen with Hrg-β1 treatment (20 ng/ml). (C) Fluorescence images of PDX#2- and PDX#3-generated tumoroids embedded in collagen, used for masking the cells for the displacement analysis in Fig. 6C-D. Scale bars, 100 μm (main).

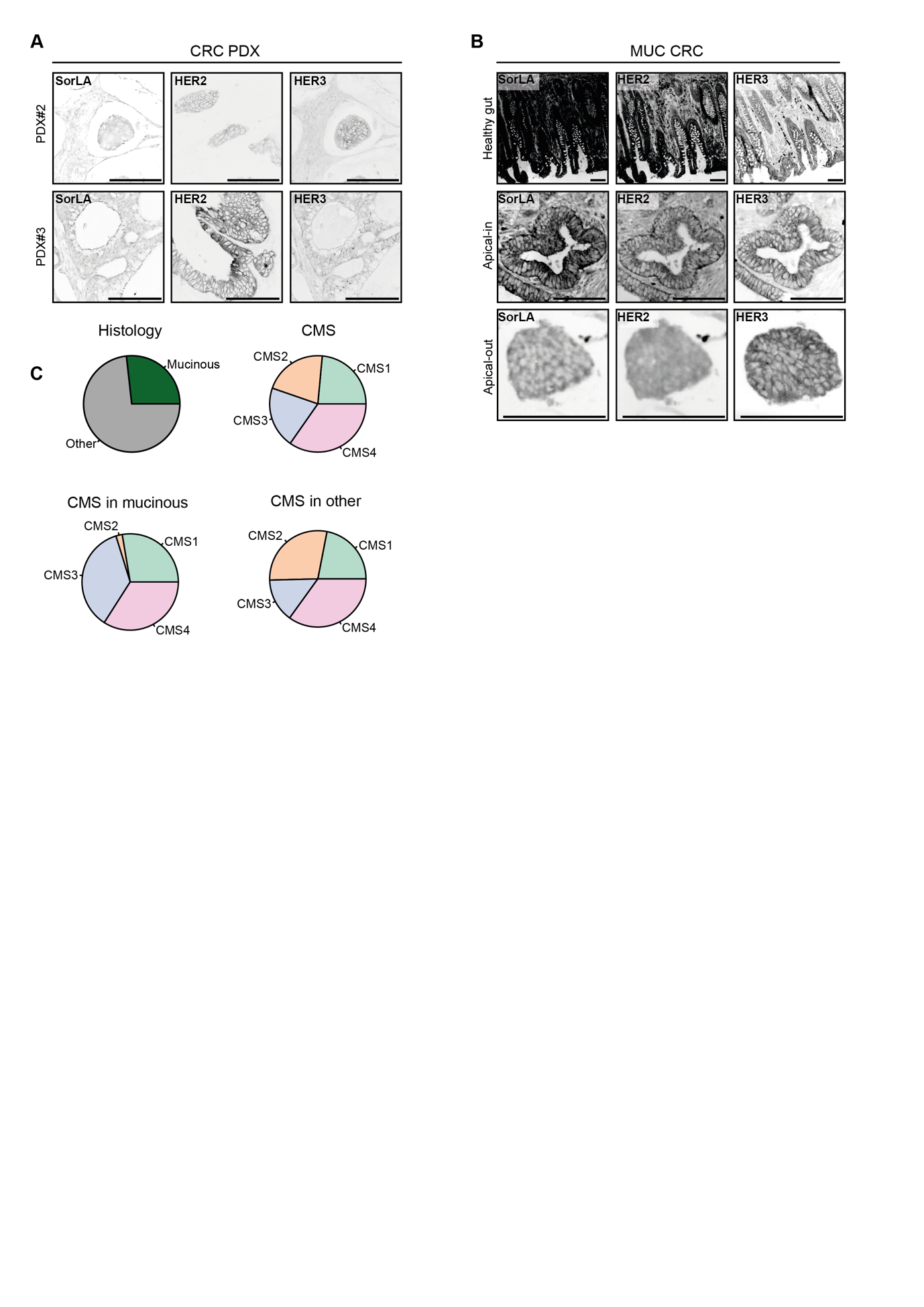

**FIGS7. HER2, HER3 and SorLA expressions are correlated *in vivo***

(A) Immunohistochemistry of PDX#2 and PDX#3 tumors stained for HER2 and SorLA, and HER2 and HER3. (B) Immunohistochemistry of healthy gut tissue, apical-in and apical-out cancer structures stained for HER2, HER3 and SorLA. Same specimens as in Fig. 7A. (C) Classification of the histotypes of the cohort used for FIG7D-E, further stratified into mucinous and non-mucinous CRCs. Scale bars, 100 μm.
