## Supplementary figures and images for "Signaling downstream of tumor-stroma interaction regulates mucinous colorectal adenocarcinoma apicobasal polarity"

### Video 1

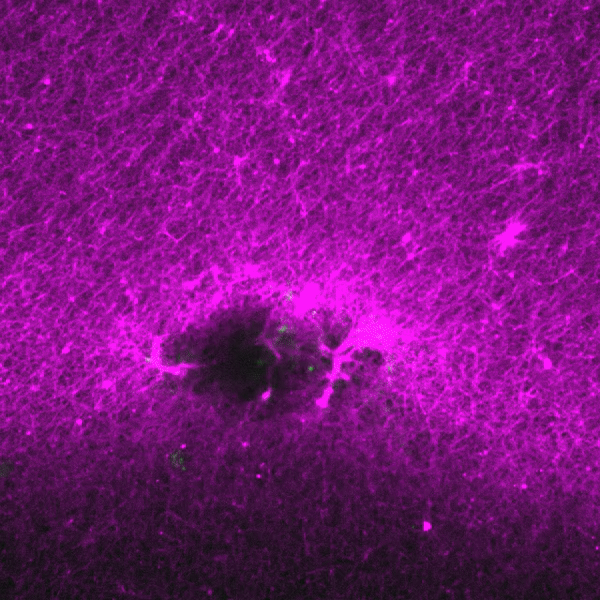

### Video 2

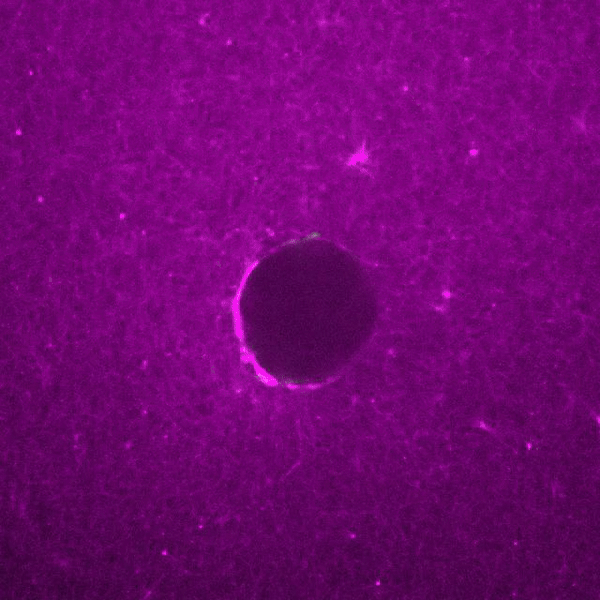
